# DNA damage and somatic mutations in mammalian cells after irradiation with a nail polish dryer

**DOI:** 10.1101/2021.02.03.429605

**Authors:** Maria Zhivagui, Areebah Hoda, Noelia Valenzuela, Yi-Yu Yeh, Jason Dai, Yudou He, Shuvro P. Nandi, Burcak Otlu, Bennett Van Houten, Ludmil B. Alexandrov

## Abstract

Ultraviolet A light is commonly emitted by UV-nail polish dryers with recent reports suggesting that long-term use may increase the risk for developing skin cancer. However, no experimental evaluation has been conducted to reveal the effect of radiation emitted by UV-nail polish dryers on mammalian cells. Here, we show that irradiation by a UV-nail polish dryer causes high levels of reactive oxygen species, consistent with 8-oxo-7,8-dihydroguanine damage and mitochondrial dysfunction. Analysis of somatic mutations reveals a dose-dependent increase of C:G>A:T substitutions in irradiated samples with mutagenic patterns similar to mutational signatures attributed to reactive oxygen species. In summary, this study demonstrates that radiation emitted by UV-nail polish dryers can both damage DNA and permanently engrave mutations on the genomes of mammalian cells. These results have far-reaching implications in regard to public health and to preventing skin cancer due to consumer-based exposure to ultraviolet light from artificial sources.

## INTRODUCTION

Ultraviolet (UV) light is a type of electromagnetic radiation with a wavelength ranging between 10nm and 400nm. Since wavelengths below 280nm are generally blocked by the Earth’s stratospheric ozone, the UV light that reaches the Earth’s surface is between 280nm and 400nm^1^. The UV spectrum can be further categorized based on its effect on human skin as well as on its ability to induce DNA damage^2^. Ultraviolet B light (UVB; 280-315nm) accounts for about 10% of the UV found on Earth, penetrates the outer layer of the skin, and it induces a plethora of DNA lesions including cyclobutane-pyrimidine dimers and 6-4 photoproducts^3,4^. In contrast, ultraviolet A light (UVA; 315-400nm) constitutes approximately 90% of the ultraviolet radiation that reaches the surface of the Earth, it can penetrate the skin more deeply, and it causes little direct DNA damage as UVA is poorly absorbed by DNA^2,5-12^.

The International Agency for Research on Cancer has classified broadband UVA (315-400nm) as a Group 1 carcinogen, based on sufficient evidence for carcinogenicity in both humans and experimental models combined with strong mechanistic considerations^13^. While UVA is found in sunlight, most of UVA environmental toxicity has been attributed to the use of commercial products, such as tanning beds^14^. Consistently, meta-analyses have shown a causal relation between skin cancer and irradiation with UV-emitting tanning devices^13,15^. Further, it has been demonstrated that skin squamous-cell carcinoma will develop in mice after long-term exposure to broadband UVA^16-19^. Prior experimental studies have also suggested that UVA irradiation leads to indirect DNA damage mostly through the accumulation of 8-oxo-7,8-dihydro-2′-deoxyguanosine (8-oxo-dG) derived from reactive oxygen species^20-24^. Studies using reporter single-gene assays have identified an enrichment of C:G>A:T mutations in UVA irradiated samples consistent with damage due to 8-oxo-dG^25-28^. Despite prior evidence for carcinogenicity of broadband UVA (315-400nm), UVA radiation in subsets of this spectrum is widely used in a surfeit of consumer products without extensive evaluation of the potential carcinogenic and mutagenic effects of these products. One prominent example is UV-nail polish dryers, which have become increasingly popular in the last decade^29,30^.

UV-nail lamps are used to cure and dry nail polish formulas, known as gels, which are oligomers requiring exposure to UV radiation to harden into polymers. These UV-gel nail devices release UVA radiation with either very little or, in most cases, no UVB radiation^31^. UV-gel nail devices contain multiple bulbs, emitting UV wavelengths between 340 and 395nm that can react and activate the photo-initiators in a gel^29^. Concerns have been raised regarding the magnitude of DNA damage that can be posed from exposure to UV-nail machines and their potential role in skin carcinogenesis^30,32^. Notably, in most cases, both nails and hands are irradiated up to 10 minutes with a UV-nail dryer per session^32^. The number of nail salon clients is estimated to reach 8 clients a day per nail technician, accounting for approximately 3 million daily clients in the United States^33^. Typically, regular users change their gel manicures every 2 weeks^32^. Recently, a small number of melanoma and non-melanoma cases, reported either on the nail or on the dorsum of the hand, have also been putatively attributed to exposure to UV radiation emitted by nail polish dryers^32,34^.

In this work, we perform an *in vitro* irradiation of human and murine cells using distinct acute and chronic exposure protocols to evaluate the DNA damage and mutagenic effects of ultraviolet radiation emitted by a nail polish UV-dryer. Irradiated and control cells are subjected to multiple assays for measuring DNA damage as well as to DNA sequencing either by bulk whole-genome sequencing after clonal expansion or by single-molecule duplex sequencing without clonal expansion (**Figure 1**). Elevated levels of reactive oxygen species (ROS) are observed in UV-irradiated samples, consistent with 8-oxo-dG damage and mitochondrial dysfunction. Analysis of somatic mutations reveals a dose-dependent increase of C:G>A:T mutations in irradiated samples with patterns similar to the ones of COSMIC signatures SBS18/36, which were previously attributed to oxidative damage^35-37^. Finally, re-examination of previously generated skin cancers uncovers that SBS18/36 are ubiquitously present in melanoma and account for ∼12% of the previously annotated driver mutations. Our results provide a comprehensive profile of the DNA damage and somatic mutations in mammalian cells after irradiation with a nail polish dryer.

**Figure 1.**
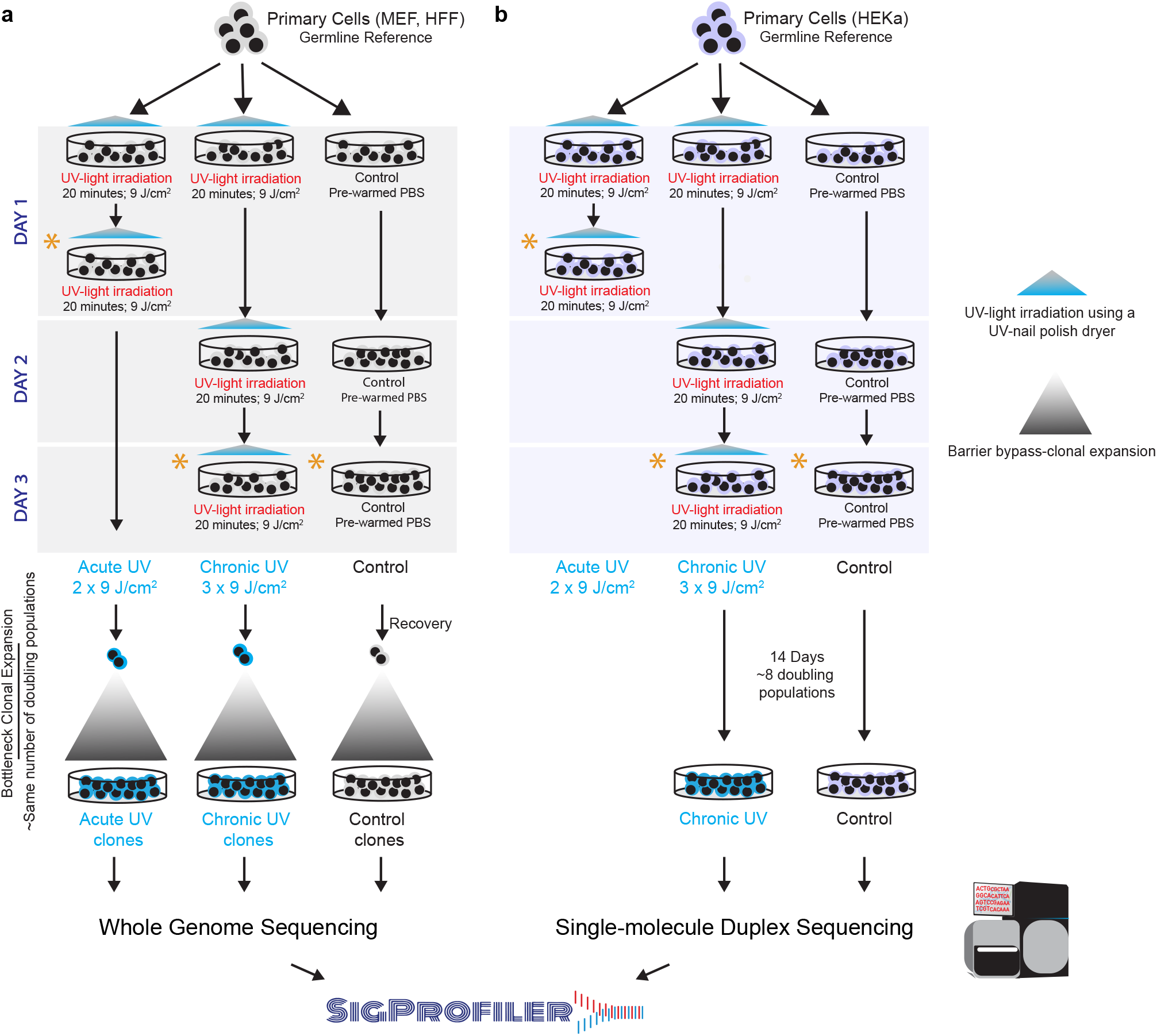
Overview of the overall study design. ***(a)*** Primary mammalian cells, MEFs and HFFs, were expanded into 6-well plates and treated with ultraviolet light (UV) emitted from a UV-nail polish dryer for 20 minutes, twice a day within one single day, termed *acute UV exposure*. For *chronic UV exposure*, primary cells were exposed consecutively in three different days with each exposure lasting 20 minutes. Control samples were maintained in the dark in pre-warmed PBS for 20 minutes during each exposure session. After recovery and cellular selection, whether through senescence bypass or single-cell subcloning, control and irradiated cell were grown for the same number of doubling populations and subjected to bulk whole-genome sequencing. Analysis of whole-genome sequenced samples was performed using our established pipelines for mutation calling. Analysis of mutational signatures was performed using the SigProfiler suite of tools. ***(b)*** Primary HEKa cells were irradiated in 10-cm dishes using UV-nail polish dryer for 20 minutes for three consecutive days. Control samples were maintained in the dark in pre-warmed PBS for 20 minutes during each exposure session. HEKa cells were kept in culture for 14 days for recovery and replication, allowing around 8 doubling-populations after which the cells were subjected to high coverage duplex sequencing. Analysis of duplex sequencing samples was performed using commercially established pipelines. Analysis of mutational signatures was performed using the SigProfiler suite of tools. Asterisks in panels *(a)* and *(b)* denote timepoints when DNA damage and other assessments were performed for each condition, including interrogation of cytotoxicity, genotoxicity, oxidative damage, and mitochondrial damage (**Supplementary Table 1**).

## RESULTS

### Experimental Design and Examination of Cytotoxicity

To study the cytotoxic effect of irradiation by a UV-nail polish dryer, mouse embryonic fibroblasts (MEFs), human foreskin fibroblasts (HFFs), and adult human epidermal keratinocytes (HEKa) were exposed under several distinct conditions (**Figure 1**). Each primary cell line was irradiated one, two, or three times, with the duration of each exposure lasting between 0 and 20 minutes. Cell viability was measured 48 hours after the final irradiation with each condition repeated at least three times. Analysis of cell viability revealed that UV radiation induced cytotoxicity with higher number of exposures causing a lower cell viability (**Figure 2*a***). For example, in all cell line models, a single 20-minute irradiation resulted in 20% to 30% cell death, while three consecutive 20-minute exposures caused between 65% and 70% cell death (q-values < 0.05; Mann-Whitney U tests; **Figure 2*a***).

**Figure 2.**
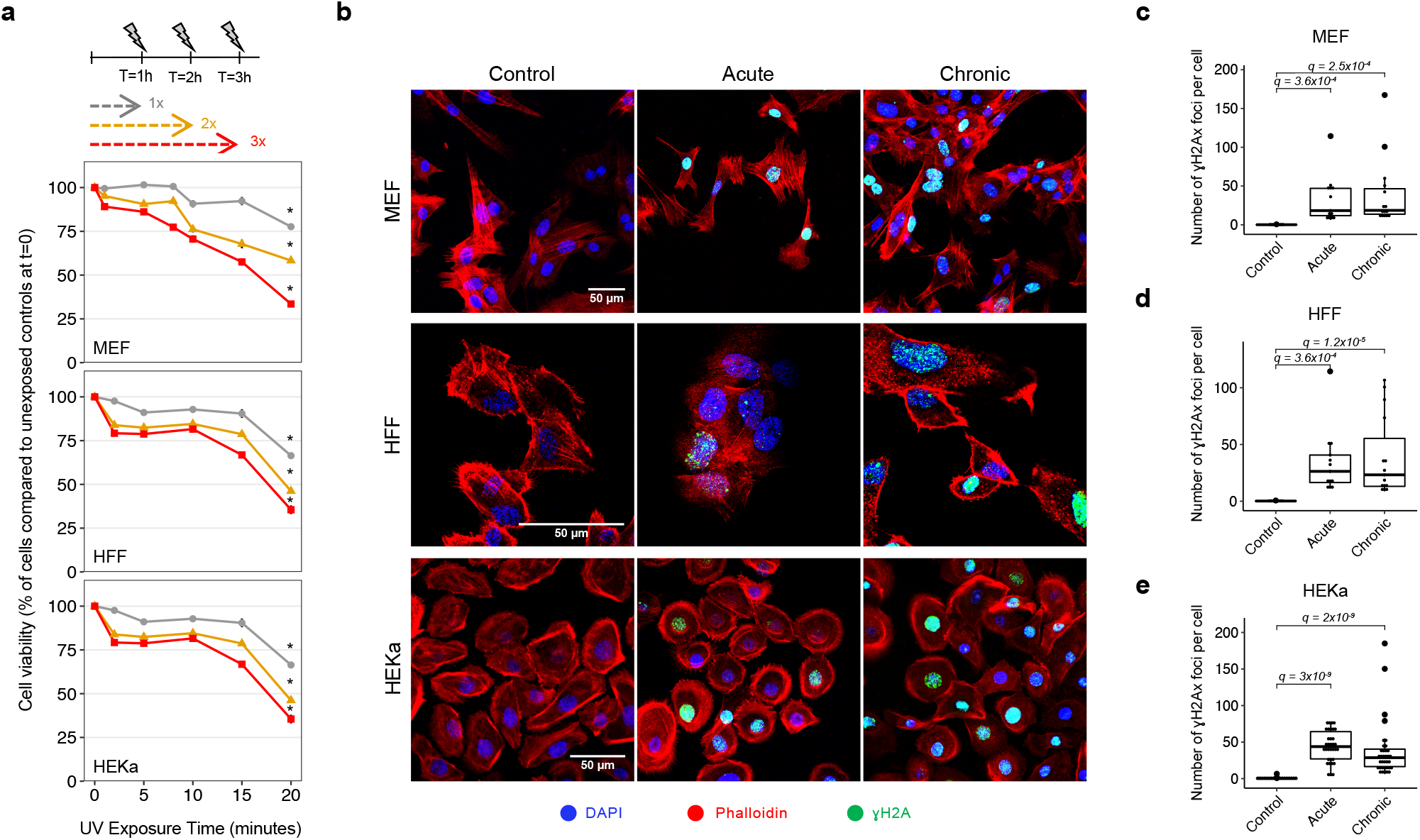
Cytotoxicity and genotoxicity in mammalian cells after irradiation with a UV-nail dryer. ***(a)*** Cytotoxicity assessment following exposure of primary MEFs (top panel), HFFs (middle panel), and HEKa (bottom panel) to UV radiation emitted from a UV-nail polish dryer for different timepoints, ranging from 0 to 20 minutes. Multiple UV-exposure sessions were tested with one hour difference between each consecutive exposure, including: grey – one exposure in a day; yellow – two exposures in a day; red – three exposures in a day. Formazan dye absorbance was measured 48 hours after treatment cessation and was normalized to the number of unirradiated control cells at timepoint 0. The results are presented as mean percentage ± standard deviation from at least four replicates (*n*=4 for MEFs, *n*=5 for HFFs, *n*=4 for HEKa). Statistically significant results from FDR corrected Mann-Whitney U two-sided tests are denoted as: *q-value<0.05. ***(b)*** DNA damage evaluation by immunofluorescence of Ser139-pjosphorylated histone H2Ax (γH2Ax). Primary MEFs (top panel), HFFs (middle panel) and HEKa (bottom panel), were exposed to UV radiation either acutely or chronically. DAPI is shown in blue, Phalloidin in red, and γH2Ax in green. ***(c-e)*** Quantification of the number of γH2Ax foci was performed by analyzing 100 cells per condition, collected from at least 3 independent experiments, in MEF (*n*=4) ***(c)***, HFF (*n*=3) ***(d)*** and HEKa (*n*=3) ***(e)*** cells. The bounds of the boxplots represent the interquartile range divided by the median, and Tukey-style whiskers extend to a maximum of 1.5 × interquartile range beyond the box. Statistically significant results from FDR corrected Mann-Whitney U two-sided tests are denoted as *q*-values.

Assessment of DNA damage was performed on MEFs and HFFs after either two 20-minute irradiations taking place within 2 hours in a single day (termed, acute exposure; **Figure 1*a***; **Supplementary Table 1**) or after three 20-minute irradiations each occurring in 3 consecutive days (termed, chronic exposure; **Figure 1*a***; **Supplementary Table 1**). Assessment of DNA damage was also performed at the same timepoint for unexposed MEF and HFF cells. Moreover, to assess the presence of somatic mutations, both irradiated and control cells were subjected to barrier bypass-clonal expansion^38^, grown for the same number of doubling populations, and subjected to bulk whole-genome sequencing (**Supplementary Table 1**; **Supplementary Figure 1**). Specifically, irradiated and control MEF cells were grown for ∼28 doubling populations prior to bulk whole-genome sequencing, while exposed and unexposed HFF cells were grown for ∼35 rounds of cell division (**Supplementary Table 1**; **Supplementary Figure 1**).

In the case of HEKa, cells were only chronically exposed with three 20-minute irradiations each occurring in 3 consecutive days (**Figure 1*b***). Assessment of DNA damage in HFFs was conducted similarly to the assessments perform for MEFs and HFFs (**Supplementary Table 1**). Nevertheless, somatic mutations in HEKa cells were detected using an orthogonal approach to the one applied for MEFs and HFFs by avoiding barrier bypass-clonal expansion. Specifically, irradiated and control HEKa cells were grown for exactly 14 days (**Figure 1*b***), corresponding to approximately 8 doubling populations, and were subsequently subjected to targeted single-molecule duplex sequencing^39^.

### Quantification of Genotoxicity

Genotoxicity of mammalian cells irradiated with a UV-nail polish dryer was evaluated 4 hours after exposure using γH2Ax immunofluorescence^40^. Statistically significant increases in the number of γH2Ax foci were observed when comparing irradiated MEF, HFF, and HEKa to control cells for all cell lines as well as for all types of exposure (**Figure 2*b-e***). For MEFs, the number of γH2Ax foci increased 186-fold for acute exposure when compared to control samples (q-value: 3.6 × 10^−4^; Mann-Whitney U test) and 183.3-fold for chronic exposure (q-value: 2.5 × 10^−4^; **Figure 2*c***). Acutely exposed HFFs had 154.7-fold elevation of the number of γH2Ax foci when compared to control samples (q-value: 3.6 × 10^−4^), while chronically exposed HFFs exhibited 137-fold elevation (q-value: 1.2 × 10^−5^; **Figure 2*d***). Similarly, irradiated HEKa cells showed a significant increase of γH2Ax foci when compared to control samples with 398-fold elevation for acute exposure (q-value: 3 × 10^−9^) and 260-fold for chronic exposure (q-value: 2 × 10^−9^; **Figure 2*e***).

### Evaluation of Photoproducts’ Formation

Immunofluorescence was used to evaluate whether irradiation from a UV-nail dryer resulted in the formation of cyclobutane pyrimidine dimers (CPDs) or pyrimidine-pyrimidone (6-4) photoproduct (6-4PPs) in all cell lines models after either acute or chronic exposure (**Supplementary Figure 2**). As a positive control, each cell line model was also exposed to ultraviolet C light (UVC), which is known to form these photoproducts^41^. Our results demonstrated that, in contrast to UVC radiation, UV emitted by a nail-polish dryer induced neither CPDs nor 6-4PPs in any of the exposed cell lines (**Supplementary Figure 2**).

### Quantification of Oxidative Damage

To evaluate oxidative damage after irradiation, we utilized three independent *in vitro* assays, including: *(i)* CellROX for detection of oxidative stress^42^; *(ii)* enzyme-linked immunosorbent assay (ELISA) for quantification of 8-oxo-dG^43,44^; and *(iii)* OxiSelect™ for detection of cytosolic and extra-cellular reactive oxygen species (ROS)^45^.

CellROX green is a fluorogenic probe used for measuring oxidative stress in live cells, namely superoxide radical anion (O_2_^•^) and hydroxyl radical (•OH)^42^. The reduced state of the CellROX dye is weakly fluorescent inside cells but once oxidized by ROS, the dye binds to nucleic acids in cells and exhibits bright green photostable fluorescence^42^. The three cell models (MEF, HFF, and HEKa) were tested using CellROX green dye immediately after irradiation (**Supplementary Table 1**). Live cell imaging results unveiled a clear increase of CellROX green fluorescence in irradiated cells compared to controls in all cases (**Figure 3*a-b***). To ensure that the CellROX fluorescence is due to ROS formation in the cells, we further challenged the HEKa cells with a ROS scavenger, namely N-acetyl-L-cysteine (NAC), prior to irradiation^46^. Quantification of the number of CellROX foci without NAC treatment revealed a significant increase of oxidative stress in irradiated HEKa cells when compared to the controls (5.5-fold increase for acute, q-value < 2.2 × 10^−16^; 4.4-fold increase for chronic, q-value < 2.2 × 10^−16^; Mann-Whitney U tests). Importantly, the number of CellROX foci significantly decreased after NAC treatment when compared to untreated cells (acute: 5.2-fold decrease, q-value= 2 × 10^−3^; chronic: 4.2-fold decrease, q-value= 1.4 × 10^−3^), confirming that irradiation by a nail polish dryer is responsible for oxidative stress induction in mammalian cells.

**Figure 3.**
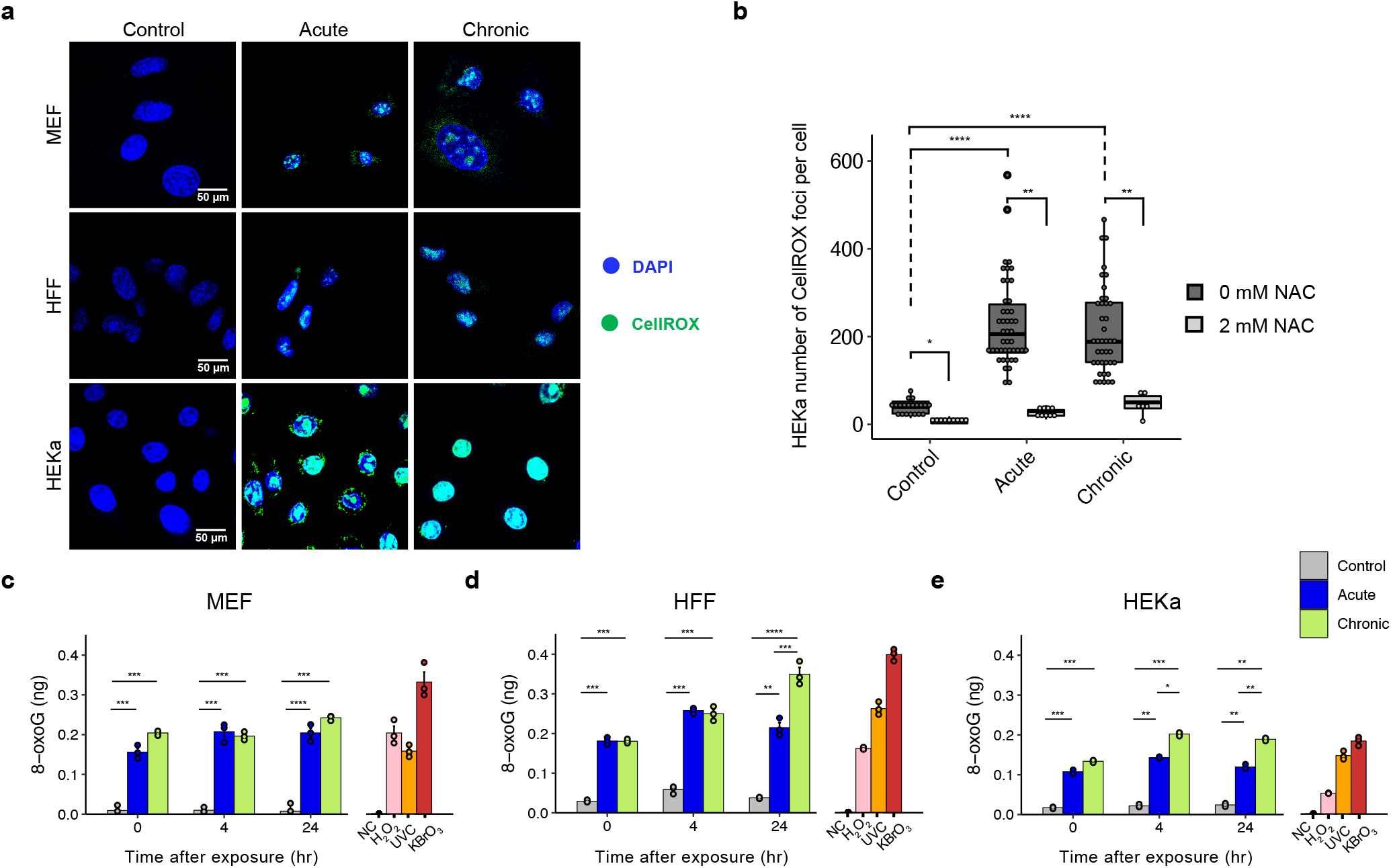
Oxidative damage in mammalian cells after irradiation with a UV-nail dryer. **(*a*)** Assessment of oxidative DNA damage by live imaging of CellROX green reagent in control and irradiated MEF (top panel), HFF (middle panel), and HEKa (bottom panel) cells. DAPI is shown in blue while CellROX is shown in green. ***(b)*** HEKa cells were challenged with 2mM N-acetyl-L-cysteine (NAC) for 1 hour prior to irradiation. Quantification of the number of CellROX foci was performed by analyzing at least 100 cells per condition and comparing cells with and without NAC pre-treatment, collected from triplicates. The bounds of the boxplots represent the interquartile range divided by the median, and Tukey-style whiskers extend to a maximum of 1.5 × interquartile range beyond the box. ***(c-e)*** Oxidative DNA damage measurement of 8-oxo-7,8-dihydro-2′-deoxyguanosine (8-oxo-dG) by ELISA in MEF ***(c)***, HFF ***(d)***, and HEKa ***(e)*** cells. Measurements of 8-oxo-dG were performed at different timepoints (0, 4, 24 hours post-treatment). Measurements of 8-oxo-dG were also performed on a negative control (NC), consisting of water only, and on three positive controls, namely, hydrogen peroxide (H_2_O_2_), ultraviolet C radiation (UVC), and potassium bromate (KBrO_3_). In all cases, data are presented as a mean value ± standard deviation from *n=3* independent biological replicates. For all panels: statistically significant results from FDR corrected Mann-Whitney U two-sided tests are denoted as: *q-value<0.05; **q-value<0.01; ***q-value<0.001; and ****q-value<0.0001.

ELISA quantifications of 8-oxo-dG were performed for all cell line modes in 3 distinct timepoints: *(i)* immediately after exposure; *(ii)* 4 hours post-irradiation; and *(iii)* 24 hours post-irradiation (**Figure 3*c-e***). Additionally, three positive controls, namely, hydrogen peroxide (H_2_O_2_), ultraviolet C radiation (UVC), and potassium bromate (KBrO_3_), were also evaluated for each of the cell lines immediately after exposure (**Figure 3*c-e***). Overall, statistically significant enrichments of 8-oxo-dG concentrations, ranging from 5-to 20-fold (q-values < 0.05), were observed in irradiated cells when compared to the control cells for all cell line models and for all exposure types (**Figure 3*c-e***). In all cases, this level of enrichment was comparable or higher to the one exhibited by the exposure to positive controls. Moreover, the amount of 8-oxo-dG remained stable even 24 hours after irradiation (**Figure 3*c-e***). These results confirm that irradiation by a UV-nail polish dryer causes high levels of oxidative stress mediated by ROS formation and subsequent oxidation of DNA bases via 8-oxo-dG formation.

To further evaluate the total free radicals in the irradiated MEF and HFF cells, we utilized OxiSelect™ *In Vitro* ROS/RNS Assay Kit. This assay employs a proprietary quenched fluorogenic probe, dichlorodihydrofluorescin DiOxyQ (DCFH-DiOxyQ), which is newly developed derivative of DCFDA and has been shown to be more specific for the detection of ROS/RNS^45,47,48^. Cytosolic and extra-cellular ROS signals were evaluated in MEF and HFF samples, in three different timepoints: *(i)* immediately after exposure, *(ii)* 20 minutes post-irradiation, and *(iii)* 24 hours post-irradiation. (**Supplementary Figure 3**). ROS-induced florescence was elevated in both acutely and chronically exposed mammalian cells compared to the untreated cells. For both MEFs and HFFs, acute exposure resulted in 2-to 8-fold increase of both cytosolic and extra-cellular ROS compared to their respective controls (q-values: 2 × 10^−5^ and 1.7 × 10^−6^ for MEF cytosolic and extracellular, respectively; q-values: 2 × 10^−3^ and 3 × 10^−5^ for HFF cytosolic and extracellular, respectively; Mann-Whitney U tests; **Supplementary Figure 3**).

### Examining Mitochondrial Membrane Potential and ROS Generation

As mitochondria are particularly sensitive to excessive ROS generated by UV radiation and they play key role in oxidants production^49,50^, we also examined mitochondrial potential membrane and evaluated the mitochondrial ROS production after irradiation with a UV-nail polish dryer. Mitochondrial activity and functionality were conducted in all three cellular models 24 hours post-irradiation by using a tetramethylrhodamine dye (TMRM), which is taken up by mitochondria with a high proton gradient and is a measure of well coupled mitochondria (**Figure 4*a***; **Supplementary Figure 4**). In all three cellular models, irradiation resulted in reduced TMRM fluorescence indicating a likely mitochondrial loss of mitochondrial membrane potential (**Figure 4*a***; **Supplementary Figure 4**). Moreover, MitoSOX red mitochondrial superoxide indicator dye was applied to HEKa cells 24 hours post-radiation. MitoSOX selectively targets the mitochondria and, after being oxidized by superoxide, it produces red fluorescence upon binding to nucleic acid in the cells. Live cell imaging of acutely and chronically irradiated HEKa cells revealed MitoSOX red fluorescence when compared to unexposed HEKa controls (**Figure 4*b***). Quantification of the number of MitoSOX foci per cell revealed a significant increase of oxidative stress in mitochondria of irradiated HEKa cells when compared to the controls (8.3-fold increase for acute, q-value: 7.4 × 10^−12^; 17-fold increase for chronic, q-value < 3.6 × 10^−10^; Mann-Whitney U tests; **Figure 4*c***). This increase in MitoSOX fluorescence putatively suggests an elevation of oxidative stress in mitochondria caused by irradiation. In principle, some forms of ROS intermediates cannot persist in the cell and the observed increase of oxidative stress in the mitochondria could be due to cellular damage leading to chronic and persistent state of hydrogen peroxide production that may cause oxidation of nuclear proteins over a 24-hour period^51^. Taken altogether, these data suggest that UVA light can cause mitochondrially generated ROS and persistent nuclear DNA damage. Nevertheless, other explanations of the observed increase of oxidative stress in the mitochondria are also possible.

**Figure 4.**
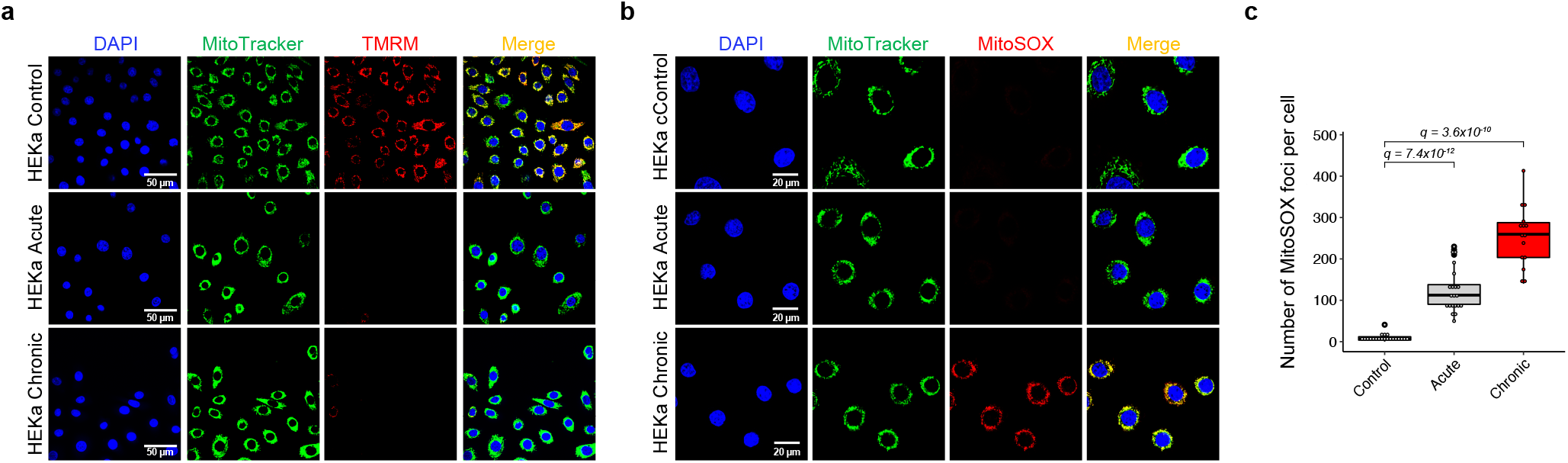
Mitochondrial disruption after irradiation with a UV-nail dryer in HEKa cells. ***(a)*** Evaluating the mitochondrial membrane potential after irradiation of HEKa cells with a UV-nail dryer. DAPI is shown in blue, MitoTracker in green, and tetramethylrhodamine dye (TMRM) in red. Yellow corresponds to overlaps between MitoTracker and TMRM. Images collected from three independent experiments. ***(b)*** Examining mitochondrial ROS production in HEKa cells. DAPI is shown in blue, MitoTracker in green, and MitoSOX in red. Yellow corresponds to overlaps between MitoTracker and MitoSOX. In both panels (*a*) and (*b*), MitoTracker green reagent is used to localize mitochondria regardless of mitochondria membrane potential. Images collected from three independent experiments. **(*c*)** Quantification of the number of MitoSOX foci in 100 cells per condition in HEKa, examined over 4 independent replicates. The bounds of the boxplots represent the interquartile range divided by the median, and Tukey-style whiskers extend to a maximum of 1.5 × interquartile range beyond the box. Statistically significant results from FDR corrected Mann-Whitney U two-sided tests are denoted as *q*-values.

### Somatic Mutations After Barrier Bypass-clonal Expansion

To evaluate the somatic mutations engraved by UV radiation from a nail dryer, we utilized two orthogonal sequencing approach: *(i)* bulk whole-genome sequencing of immortalized HFF and MEF clones, grown for the same number of doubling populations after barrier bypass-clonal expansion (**Supplementary Figure 1**); and *(ii)* single molecule duplex sequencing for HEKa cells without clonal expansion (**Figure 1*b***). A total of 15 HFF and 15 MEF immortalized clones were subjected to bulk whole-genome sequencing (WGS) at 30x coverage, including: *(i)* 5 HFF and 5 MEF clones after acute irradiation; *(ii)* 5 HFF and 5 MEF clones after chronic irradiation; *(iii)* 5 HFF and 5 MEF unirradiated control clones. Somatic mutations were derived for immortalized clones by comparing the WGS data of each immortalized clone to the WGS data from the respective primary cell line using previously established state-of-the-art bioinformatics pipelines (**Figure 1**; **Supplementary Figure 5**). Examining the variant allele frequencies (VAFs) for single base substitutions revealed that the majority of MEF mutations are subclonal with a mean VAF of approximately 0.25, while most HFF mutations are clonal with a mean VAF of approximately 0.50 (**Supplementary Figure 6*a-b***). These results are consistent with previous observations in these model systems^38,52^ and they reflect experimental differences between the murine and human cell line models after clonal expansion. Specifically, MEFs undergoing barrier-bypass clonal expansion are most likely to yield two subclonal populations^38,52^, while HFFs undergoing single-cell subcloning have a high propensity for selecting a single clone^38^.

Single base substitutions (SBSs) were 2- and 2.5-fold elevated in acutely and chronically irradiate MEF cells, respectively, when compared to control cells (q-values<0.01; Mann-Whitney U tests; **Figure 5*a***). Stratification of SBSs based on their simplest six channel classification (*i*.*e*., C>A, C>G, C>T, T>A, T>C, T>G; each mutation referred to by the pyrimidine base of the Watson-Crick DNA base-pair) revealed a significant increase of C>A transversions in acute and chronic MEFs exposed to UV emitted from a nail dryer: 2.8- and 4.2-fold increase for acute and chronic, respectively (q-values: 0.02 and 0.016; Mann-Whitney U tests; **Figure 5*b***). Importantly, the number of C>A single base substitutions found in MEF clones was positively correlated with the number of UV exposures (**Figure 5*c***; Spearman’s rank correlation: 0.64; p*-*value: 0.0098).

**Figure 5.**
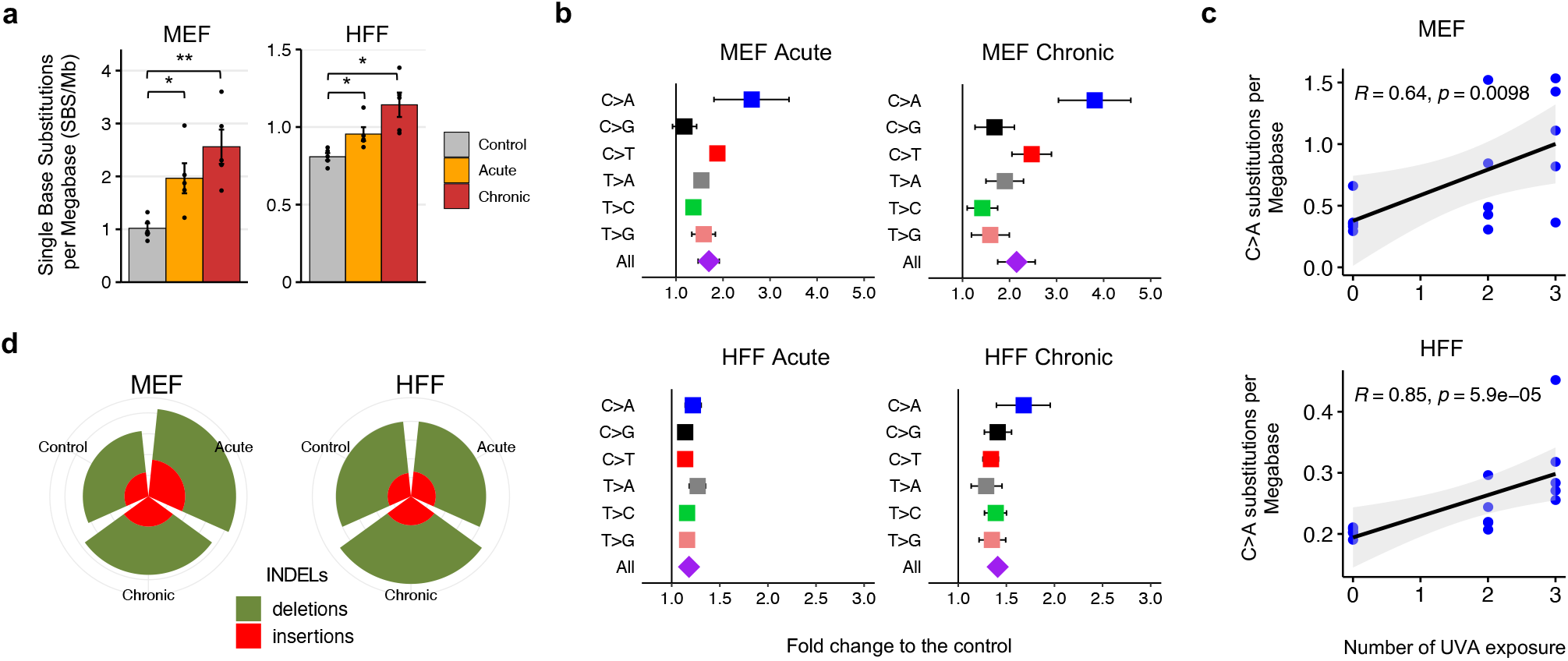
Mutations found in the genomes of MEFs and HFFs irradiated by a UV-nail dryer. ***(a)*** Mutation count per megabase (Mb) detected in the different conditions, represented in colors, in MEFs and HFFs. Data is presented as a mean value ± standard error from *n=5* independent biological replicates. Statistically significant results from FDR corrected Mann-Whitney U two-sided tests are denoted as: *q-value<0.05; **q-value<0.01. ***(b)*** Fold-increase of single base substitutions in UV-treated clones compared to controls. Fold increase is expressed as mean fold-change ± SE (standard error) from *n=5* independent replicates. ***(c)*** Spearman’s rank correlations between the number of C>A substitutions and the number UV exposures in the five independent UV-treated clones in MEFs (top panel) and HFFs (bottom panel). Acute and chronic exposures correspond to 2 and 3 exposures, respectively. P-value is calculated based on a two-sided t-distribution with n-2 degrees of freedom. Grey bands represent the 95% confidence intervals. ***(d)*** Number of small insertions and deletions (indels) in MEF and HFF clones for each irradiation condition.

Immortalized HFF clones exhibited similar behaviors to the ones of MEF clones, albeit, in most cases, with smaller effect sizes. Specifically, substitutions were 1.2- and 1.4-fold enriched in acutely and chronically irradiated HFF cells compared to control cells (q-values<0.01; Mann-Whitney U tests; **Figure 5*a***). An elevation of C>A mutations was also observed in acutely irradiated HFF clones (1.22-fold increase with q-value: 0.015; Mann-Whitney U tests; **Figure 5*b***) as well as in chronically irradiated HFF clones (1.63-fold increase with q-value: 7.9 × 10^−3^; Mann-Whitney U tests; **Figure 5*b***). Similarly, the number of C>A transversions exhibited a positive correlation with the number of UV exposures in HFFs (**Figure 5*c***; Spearman’s rank correlation: 0.85; p*-*value: 5.9 × 10^−5^).

Analysis of somatic copy number changes revealed that all HFF samples are almost perfectly diploid with the exception of one chronically exposed HFF sample, which showed loss of heterozygosity in several large chromosomes. Consistent with prior reports^38,52^, focal copy-number changes were found in some of the spontaneous and irradiated MEF samples, which may lead to an inaccurate estimation of the mutational burden caused by genomic instability in MEFs. Specifically, four of the MEF samples were found to have high number of somatic copy-number changes with evidence for potential whole-genome doubling followed by loss of heterozygosity. These four MEF samples included: one spontaneous, one acutely irradiated, and two chronically irradiated clones. Nevertheless, no statistically significant differences in overall somatic copy number changes or in overall ploidy were observed between the irradiated and the control cells for any of the utilized cell line models. Further, no differences in the numbers of small insertions or deletions were observed between the irradiated and the control cells for any of the used cell line models (**Figure 5*d***).

### Somatic Mutations After Duplex Sequencing

In principle, detection of somatic mutations in experimental systems requires clonal expansion^38^ and, for MEF and HFF cell lines, we utilized barrier bypass-clonal expansions prior to bulk whole-genome sequencing. Recently, single-molecule duplex sequencing approaches have provided an alternative high-fidelity protocol for directly measuring the mutagenic impact of genotoxic agents without the need for clonal expansion, whilst improving the accuracy of NGS by more than 10,000-fold and enabling sensitive detection of rare mutations, such as those induced by mutagens^39,53,54^. Importantly, duplex sequencing does not sequence a single cell or even a small number of cells, rather, the method generally interrogates millions of double-stranded DNA coming from many thousands of cells^55^. Thus, the mutational profile from duplex sequencing provides an average somatic mutational profile across the complete population of exposed cells^39,53,54^.

We used a commercial targeted duplex sequencing protocol for assessing mutagenesis without clonal expansion^39^ both to provide an orthogonal validation of the previous results and to account for any potential artifacts due to clonal expansion in MEF and HFF cell lines. Specifically, we detected somatic mutations using duplex sequencing data from 3 chronically irradiated and 3 unirradiated control HEKa cells (**Method**). Similar to irradiated HFFs and MEFs, chronically exposed HEKa exhibited 1.3-fold increase of substitutions when compared to control cells (**Figure 6*a***; q-value: 0.02; Mann-Whitney test). Importantly, C>A transversions were 2.2-fold enriched in chronically irradiate HEKa cells compared to control cells (q-values: 0.04; Mann-Whitney tests; **Figure 6*b***). Moreover, a positive correlation between the number of C>A transversion and the number of UV exposures was also evident in HEKa cells (**Figure 6*c***; Spearman’s rank correlation: 0.88; p*-*value: 0.021). Consistent with prior results, no changes were observed for small insertions or deletions between irradiated and control HEKa cells (**Figure 6*d***).

**Figure 6.**
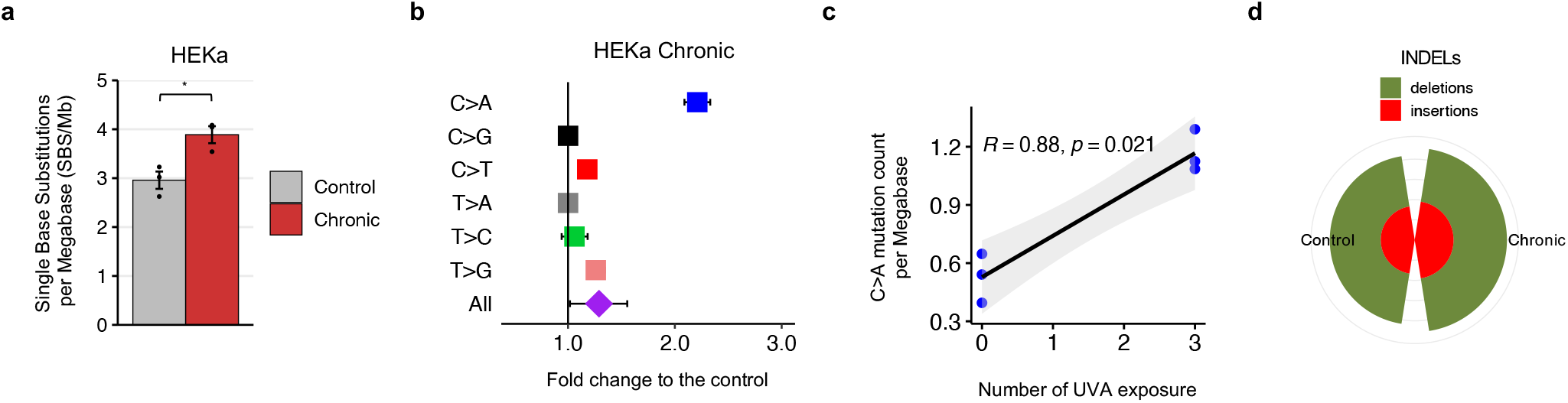
Mutation analysis of HEKa cells irradiated by a UV-nail dryer. ***(a)*** Mutation count per megabase (Mb) detected in the different conditions, represented in colors, in HEKa cells. Data is presented as a mean value ± standard error from *n=3* independent biological replicates. **(*b*)** Fold-increase of single base substitutions in UV-treated clones compared to controls. Fold increase is expressed as mean fold-change ± SE (standard error) from *n=3* independent replicates. ***(c)*** Spearman’s rank correlations between the number of C>A substitutions and the number UV exposures in each of the 3 biological replicates. P-value is calculated based on two-sided t-distribution with *n-2* degrees of freedom. Grey bands represent the 95% confidence intervals. ***(d)*** Number of small insertions and deletions (indels) in HEKa cells. Statistically significant results from FDR corrected Mann-Whitney U tests are denoted as: *q-value<0.05; **q-value<0.01; ***q-value<0.001; and ****q-value<0.0001.

### Mutational Signatures Engraved by Irradiating with a UV-nail Polish Dryer

Previously, we have shown that different endogenous and exogenous mutational processes engrave characteristic patterns of somatic mutations, termed, mutational signatures^56-58^. To evaluate the mutational signatures engraved by irradiation with a UV-nail polish dryer, we performed separate analysis of mutational signatures for MEFs, HFFs, and HEKa data. To explain the SBS patterns of mutations observed in the irradiated cells, any known COSMIC signature^59^ that improved the reconstruction above the background models was allowed. For the analyses of MEF data, in addition to known COSMIC signatures, we also allowed mouses specific mutational signatures previously observed either *in vitro*^52^ or *in vivo*^60^ murine models (**Supplementary Figure 7*a*)**.

Our analyses revealed that only COSMIC signatures SBS18 and SBS36 (SBS18/36), both previously attributed to reactive oxygen species^61,62^, were found to be operative in all irradiated MEF and HFF cells. It should be noted that ROS signatures SBS18 and SBS36 have cosine similarity of 0.914 (**Figure 7*a***), which, in many cases, makes them hard to distinguish from one another and these signatures are commonly reported together as SBS18/36^35,63,64^. Our analyses revealed that SBS18/36 were the only signatures enriched in irradiated HFF and MEF samples when compared to controls (**Figure 7*b-c***; q-values<0.05; Mann-Whitney tests). No other mutational signature found in these samples was elevated in irradiated samples when compared to control samples (**Supplementary Figure** 7***b&d***). Moreover, the identified sets of operative mutational signatures allowed accurately reconstructing the mutational patterns in both MEFs and HFFs (**Supplementary Figure** 7***c&e***). The statistically significant increase of single base substitutions between unirradiated and chronically irradiated cell lines that was attributed to the ROS-associated signatures SBS18/36 was 0.70 and 0.14 mutations per megabase for MEF and HFF, respectively (**Supplementary Figure 7**). This corresponds to an additional 2,000 and 400 ROS-associated somatic mutations engraved in the genome of a chronically irradiated MEF and HFF cell, respectively.

**Figure 7.**
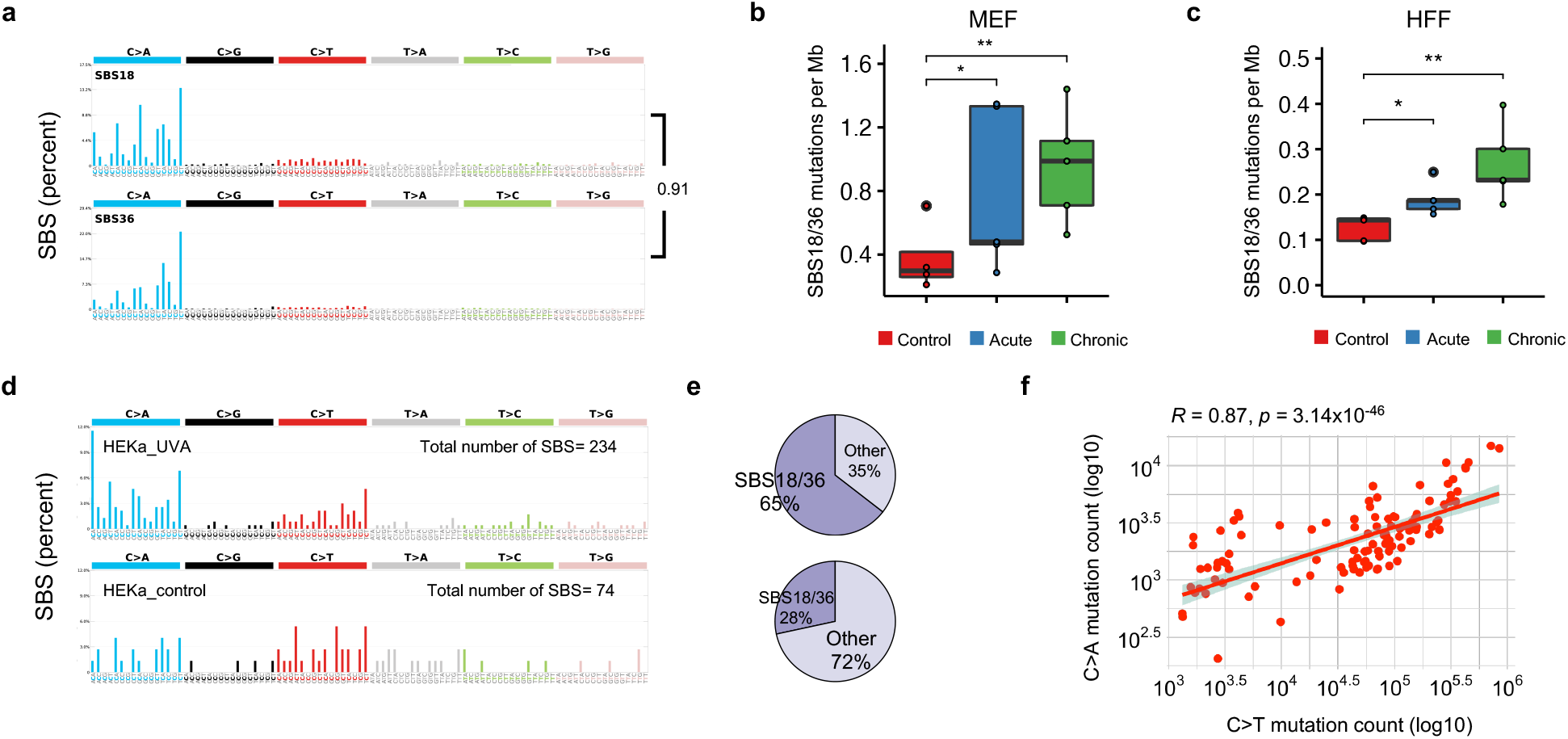
Somatic Mutations due to COSMIC signatures SBS18/36 in cells and cancer samples. ***(a)*** The patterns of mutational signatures COSMIC SBS18 and SBS36. ***(b)*** Number of mutations per megabase attributed to COSMIC SBS18/36 in UV exposed MEF clones, per condition, extracted from *n=5* biological replicates per condition. **(*c*)** Number of mutation per megabase attributed to COSMIC SBS18/36 in UV exposed HFF clones, per condition, extracted from *n=5* biological replicates per condition. **(*d*)** Mutational patterns of irradiated (upper track) and control (lower track) primary HEKa cells. **(*e*)** COSMIC signatures assignment of UV-irradiated HEKa cells (upper track) and control (lower track). **(*f*)** Pearson’s correlation of C>A and C>T mutations in *n*=*144* skin cancer samples from the PCAWG project. P-value is calculated based on two-sided t-distribution with *n-2* degrees of freedom. The bounds of the boxplots represent the interquartile range divided by the median, and Tukey-style whiskers extend to a maximum of 1.5 × interquartile range beyond the box.

Observing signatures SBS18/36, which are almost exclusively characterized by C>A substitutions (**Figure 7*a***), in irradiated clones confirms that radiation emitted by a UV-nail polish dryer not only oxidizes DNA but also results in permanently engraved patterns of C>A somatic mutations on the genomes of irradiated cells. Similarly, an almost 3-fold enrichment of signatures SBS18/36 (p-value < 0.0001; Fisher’s exact test) was observed in keratinocytes irradiated with a UV-nail polish dryer when compared to controls. Specifically, 65% of the mutations in UV-irradiated HEKa cells were attributed to SBS18/36 (**Figure 7*d-e***). In contrast, in unirradiated control samples, SBS18/36 accounted for 23% of mutations (**Figure 7*d-e***).

Re-examination of the recently published set of 144 whole-genome sequenced skin cancers from the Pan-Cancer Analysis of Whole Genomes (PCAWG) project^65^ revealed a positive correlation between C>A and C>T mutations (Pearson’s correlation: 0.87; p*-*value < 2.2 × 10^−16^; **Figure 7*f***). Moreover, an average skin cancer appears to harbor approximately 2,500 C>A substitutions (95% CI: 2,015 – 2,868) in their genome with approximately 80% of their pattern attributed to signatures SBS18/36. However, these C>A substitutions account for only 2.3% of the mutations observed in the 144 melanomas; 85% of mutations in these melanomas are generated by signature 7 – mutational signature attributed to cyclobutane-pyrimidine dimers and 6-4 photoproducts from ultraviolet light^65^. Importantly, 20 of the 169 substitutions (11.83%) identified as driver mutations^65^ in these samples can be attributed to signatures SBS18/36. This indicates that, while signatures SBS18/36 generate low numbers of mutations in skin cancer, they play an important role in generating somatic mutations that contribute towards tumor development and cancer evolution.

## DISCUSSION

In this report, we employed well-controlled experimental models for UV irradiation using a UV-nail polish dryer and evaluated the DNA damage and mutagenic changes engraved on the genomes of mammalian cells (**Figure 1**). The utilized *in vitro* clonal expansion models, MEFs and HFFs, manifest the specific features required for mimicking human carcinogenesis and for recapitulating the activity of characteristic mutational processes^38^. Importantly, the MEF model has proven to be an invaluable *in vitro* model system for emulating human mutational signatures of various exogenous and endogenous mutagens, including UVB and UVC radiation, known tobacco carcinogens (*e*.*g*., benzo[*a*]pyrene), aristolochic acid, and activation-induced cytidine deaminase (AID)^66-70^. Both MEFs and HFFs underwent clonal expansion after a barrier bypass step (senescence for MEFs and single-cell bottleneck for HFFs) leading to the formation of exposure-derived clones, which were grown for the same number of doubling populations (**Supplementary Figure 1**). Nevertheless, in some cases, these models may exhibit high background mutations due to genomic instability occurring during clonal expansion process. As an orthogonal approach, we also utilized HEKa primary cells, which cannot go through clonal expansion in culture. Cytotoxicity and genotoxicity were observed in all irradiated cells; importantly, we observed an elevation of ROS in irradiated cells leading to increased levels of 8-oxo-dG, which was confirmed by multiple *in vitro* assays (**Figure 3**). Moreover, a potential mitochondrial dysfunction was observed upon UVA radiation (**Figure 4** and **Supplementary Figure 4**). Genomic profiling revealed higher levels of somatic mutations in irradiated cells with C>A mutations being highly elevated in exposed samples (**Figures 5** and **6**). Consistent with the increase of ROS, COSMIC signatures SBS18/36, two mutational signatures previously attributed to reactive oxygen species^61,62^, were engraved on the genomes of irradiated samples and enriched in irradiated samples when compared to controls (**Figure 7**).

Broadband UVA (315-400nm) has been extensively studied in the context of tanning devices and classified as carcinogen by the International Agency for Research on Cancer^13^. Broadband UVA (315-400nm) penetrates deep in the skin, reaching the papillary dermis layer with longer-wavelength UVA even reaching the deep dermis^71^ and the stem cell compartments of the skin^72^. Prior experimental studies with broadband UVA have shown that it causes an accumulation of 8-oxo-dG^20-24^, generates C>A mutations in single-gene assays^25-28^, and even induces tumors in mice^73^. Indeed, prior studies have shown that UVA can generate low level of C>T somatic mutations consistent with pyrimidine-pyrimidine photodimers^74^. Intriguingly, in this study, we demonstrate that UVA with wavelengths between 365 and 395nm, which is generally considered to be safe and is commonly used in a plethora of consumer products, causes DNA oxidative damage leading to C>A mutations; no evidence was found that radiation generated by the UV-nail polish machine generates any pyrimidine-pyrimidine photodimers (**Supplementary Figure 2**). Importantly, the longer-wavelength of UVA emitted by a nail-polish dryer (365-395nm) will reach all layers of the epidermis and it will penetrate towards the deeper layers of the dermis, potentially, even affecting some skin stem cells. As keratinocytes are present in all layers of the epidermis and melanocytes are found in the basal layer of epidermis, exposure to UVA from a nail-polish dryer may irradiate these cells in a physiologically normal human skin.

Our findings are also consistent with prior cancer epidemiological observations. In principle, males have significantly higher risk than females to develop melanoma^75,76^. Surprisingly, this trend is reversed when one examines cancer risk in melanomas of the hand in younger women. Specifically, young females (ages 15 to 39) have approximately 2-fold higher risk for developing melanoma of the upper extremities compared to young males^77^. This higher risk is possibly due to lifestyle choices, one of which may be radiation from consumer products such as UV-nail polish dryers^77^. It should be noted that exposure to UV-light from the sun cannot reasonably explain the higher risk in young females as there is no observed differences in melanoma risk between young males and females for other parts of the body commonly exposed to the sunlight, *viz*., face, head, and neck^76,77^.

While this report demonstrates that radiation from UV-nail polish dryers is cytotoxic, genotoxic, and mutagenic, it does not provide direct evidence for increased cancer risk in human beings. Prior studies have shown that an increase in mutagenesis will likely lead to an increase in cancer risk^78-80^. Further, several anecdotal cases have demonstrated that cancers of the hand are likely due to radiation from UV-nail polish dryers^32,34^. Taken together, our experimental results and the prior evidence strongly suggest that radiation emitted by UV-nail polish dryers may cause cancers of the hand and that UV-nail polish dryers, similar to tanning beds^81^, may increase the risk of early-onset skin cancer^82^. Nevertheless, future large-scale epidemiological studies are warranted to accurately quantify the risk for skin cancer of the hand in people regularly using UV-nail polish dryers. It is likely that such studies will take at least a decade to complete and to subsequently inform the general public. In the meantime, this report aims to raise awareness of the likely cancer-causing role of artificial UV lamps and caution people who regularly use UV-nail polish dryers.

## METHODS

### UV-nail polish machine characteristics

A 54-Watt UV nail drying machine was purchased (model: MelodySusie), harboring 6 bulbs that emit UV photons for curing gel nail polish. Based on the manufacturer’s specifications, the UV nail drying machine emits ultraviolet A light in wavelengths between 365 and 395nm. The UV power density was evaluated using a UV513AB Digital UVA/UVB Meter that can measure UV radiation between 280 and 400 nm. The machine stabilizes at ∼7.5 mW/cm^2^, within minutes (**Supplementary Figure 8*a-b*)**, putting it on the lower end of the power density spectrum calculated for commonly used UV-nail machines^31^ (median of 10.6 mW/cm^2^). In this study, in most cases, we performed a 20-minute UV exposure session which is equivalent to a total amount of energy delivered per unit area of 9 J/cm^2^: *7*.*5 mW/cm*^*2*^ *\* 20 minutes = 7*.*5 mJ/(sec*cm*^*2*^*) * 1200 sec = 9000 mJ/cm*^*2*^ *= 9 J/cm*^*2*^.

### Cell culture, irradiation, and immortalization of mouse embryonic fibroblasts (MEFs)

Primary mouse embryonic fibroblasts (MEFs) were purchased from Lonza (M-FB-481). They were expanded in Advanced DMEM supplemented with 15% fetal bovine serum, 1% penicillin/streptomycin, 1% sodium pyruvate and 1% glutamine, and incubated in 20% O_2_ and 5% CO_2_. At passage 2, the primary cells were seeded in six-well plates for 24 hours, until adherence, then exposed for 20 minutes to the UV-nail machine in pre-warmed sterile PBS. Control cells were kept in PBS for 20 minutes. Cells were irradiated with a UV nail polish dryer, acutely (*i*.*e*., twice a day, with 60 minutes break between each of the 2 sessions) as well as chronically (*i*.*e*., once every day for up to 3 consecutive days). At the end of every irradiation, the cells were washed, and complete pre-warmed medium was replenished. Exposed and control primary cells were cultivated until bypassed a barrier step through senescence. Upon barrier bypass, the cells reached clonal expansion allowing the isolation of *bona fide* cell clones (**Supplementary Figure 1*a*)**. In all cases, irradiated and control MEF cells were grown for ∼28 doubling populations prior to bulk whole-genome sequencing. Fifteen clonally expanded populations were successfully isolated and subjected to bulk whole-genome sequencing, including: 5 replicates acute exposure, 5 replicates of chronic exposure, and 5 replicates for unirradiated control cells. MEFs were authenticated by PCR of the short tandem repeats (STR) to confirm the correct species and that they are contamination-free. All cell cultures were routinely tested for the absence of mycoplasma.

### Cell culture, irradiation, and immortalization of human foreskin fibroblasts (HFFs)

Primary human cells derived from human foreskin fibroblasts (HFFs) were provided to us by Dr. John Murray (Indiana University Bloomington). Early passage cells were expanded in Advanced DMEM supplemented with 10% fetal bovine serum and 1% glutamine in 5% CO_2_ incubator. Irradiation with a UV-nail polish dryer followed the same protocol as the one for MEFs (**Figure 1*a*)**. Similarly, control HFF cells were kept in PBS for 20 minutes during all irradiations. Clonal expansion was carried out using serial dilutions procedure in 96-well plates, assuming 30% probability of a single-cell clone formation per well (**Supplementary Figure 1*b*)**. Wells were washed weekly, until clones reached confluency and were transferred progressively to T-75 flasks. In all cases, irradiated and control HFF cells were grown for ∼35 rounds of cell division. Fifteen clonally expanded populations were successfully isolated and subjected to bulk whole-genome sequencing, including: 5 replicates acute exposure, 5 replicates of chronic exposure, and 5 replicates for unirradiated control cells. HFFs were authenticated by PCR of the short tandem repeats (STR) to confirm the correct species and that they are contamination-free. All cell cultures were routinely tested for the absence of mycoplasma.

### Cell culture and irradiation of adult human epidermal keratinocytes (HEKa)

Primary human keratinocytes, derived from normal adult human epidermal keratinocytes (HEKa), were purchased from ATCC (PCS-200-011; Lot number: 70033063). Early passage cells were expanded in Dermal Cell Basal Medium (PCS-200-030) supplemented with Keratinocyte Growth Kit (PCS-200-040) and antibiotics (Penicillin: 10 Units/mL, Streptomycin: 10 µg/mL, Amphotericin B: 25 ng/mL) in 5% CO_2_ incubator. At passage 2, the primary cells were seeded at a density of 2.5 × 10^3^ cells/cm^2^ in 10-cm dishes for 24 hours. Subsequently, cells were only chronically irradiated with a UV-nail polish dryer following the same protocol as the for MEFs and HFFs (**Figure 1*b*)**. Similarly, control HEKa cells were kept in PBS for 20 minutes during all irradiations. In all cases, irradiated and control HEKa cells were grown for exactly 14 days, corresponding to approximately 8 rounds of cell division, which allows sufficient time for DNA adduct/damage to be fixed as somatic mutation(s) but not enough time to result in a clonal expansion^39^. Unexposed cells were grown for the same duration as irradiated cells. Six cell populations were successfully isolated and subjected to subjected to duplex sequencing, including: 3 replicates for chronic exposure and 3 replicates for unirradiated control cells. HEKa cells were authenticated by PCR of the short tandem repeats (STR) to confirm the correct species and that they are contamination-free. All cell cultures were routinely tested for the absence of mycoplasma.

### Cell viability and cytotoxicity assays

Primary cells were seeded in 24-well plates and exposed to the UV drying device as indicated. Cell viability was measured 48 hours after treatment cessation using the Cell-Counting Kit 8 (CCK-8) from Dojindo. Plates were incubated for 3 hours at 37°C and absorbance was measured at 450 nm using the Infinite 200 Tecan *i*-control plate reader machine. The CCK-8 assay was performed in at least 4 replicates for each experimental condition. Trypan Blue exclusion assay was also performed for assessing cell viability upon exposure, validating the choice for selecting the irradiation condition that causes around 50% cell death (**Supplementary Figure 8*c*)**.

### Assessment of genotoxicity using γH2Ax immunofluorescence

Immunofluorescence staining was carried out using a monoclonal antibody specific for Ser139-phosphorylated H2Ax (γH2Ax) (9718, Cell Signaling Technology). Briefly, primary cells were seeded on coverslips in 24-well plates and, the following day, irradiated in triplicates as previously described. Four hours after treatment cessation, the cells were fixed with 4% formaldehyde at room temperature for 15 minutes, followed by blocking in 5% normal goat serum (5425, Life Technologies) for 60 minutes. Subsequently, the cells were incubated with γH2Ax-antibody (1:400 in 1% BSA) at 4°C, overnight. A fluorochrome-conjugated anti-rabbit secondary antibody (4412, Cell Signaling Technology), diluted to 1:1000, was then incubated for 1 hour at room temperature. ß-actin staining was followed using phalloidin (8953, Cell Signaling Technology) incubated for 15 minutes at room temperature. Coverslips were mounted in ProLong Gold Antifade Reagent with DAPI (8961, Cell Signaling Technology), overnight. Immunofluorescence images were captured using a Confocal Laser Scanning Biological Microscope Olympus FV1000 Fluoview. Quantification of γH2Ax fluorescent cells was computed using Fiji software (version 2.3.0).

### Investigation of photoproducts using CPD and 6-4PP immunofluorescence

Immunofluorescence staining was carried out using anti-cyclobutane pyrimidine dimers (CPDs) monoclonal antibodies (clone KTM53, Kamiya Biomedical) and anti-6-4 photoproducts monoclonal antibodies (Clone 64M-2, Cosmo Bio). Briefly, primary cells were seeded on coverslips in 24-well plates and, the following day, irradiated in triplicates as previously described. Subsequently, the cells were washed with ice-cold CSK buffer, containing 100mM NaCl, 300mM Glucose, 10mM PIPES, 3mM MgCl2 and 0.5% Triton, then fixed with 4% formaldehyde at room temperature (RT) for 15 minutes, followed by permeabilization using 0.5% Triton for 5 minutes on ice. DNase I treatment was performed to denaturalize the genomic DNA in the cells using 20U DNase I for 40s per coverslip (NEB; DNase I (RNase-free), M0303S). For blocking, we used 1% BSA (A3059, Sigma) and 5% normal goat serum (5425, Life Technologies) for 60 minutes. Afterwards, the cells were incubated with the monoclonal antibodies (1:50 in 1% BSA) at 4°C, overnight. A fluorochrome-conjugated anti-mouse Alexa Fluor 594 secondary antibody (8890S, Cell Signaling Technology), diluted to 1:1000, was then incubated for 1 hour at room temperature. Coverslips were mounted in ProLong Gold Antifade Reagent with DAPI (8961, Cell Signaling Technology), overnight. Immunofluorescence images were captured using Nikon A1R-STORM Super Resolution Microscope and processed on Fiji software (version 2.3.0).

### Quantification of reactive oxygen species via CellROX

Oxidative stress induces the production of reactive oxygen species (ROS) and reactive nitrogen species (RNS) in cells. We employed the CellROX™ Green Reagent for detection of oxidative stress (ThermoFisher Scientific; catalog number: C10444), immediately after irradiation. MEF, HFF, and HEKa primary cells were seeded on Mattek 35-mm dishes from (P35G-1.5-14-C) appropriate for high quality live cell imaging. CellROX green was added in pre-warmed complete media to a final concentration of 5µM and incubated for 30 minutes at 37 °C. CellROX solution was washed carefully 3 times with pre-warmed PBS and normal media was replenished. Live cell imaging was conducted using Nikon A1R-STORM Super Resolution Microscope and processed on Fiji software (version 2.3.0). To ensure CellROX selectivity towards ROS components, cells were challenged with 2mM N-acetyl-L-cysteine (NAC) for 1 hour prior to UVA exposure. NAC is a known ROS scavenger. After CellROX addition, live cell images were used for quantification of CellROX foci number, with and without NAC pre-treatment, using General Analysis pipeline on Nikon WorkStation 5317.

### Quantification of reactive oxygen species via OxiSelect

Additionally, we also utilized the OxiSelect™ *In Vitro* ROS/RNS Assay Kit (Green Fluorescence), from Cell Biolabs, to evaluate the level of oxidative damage induced after irradiation, intra- and extra-cellularly, immediately after exposure as well as 20 minutes and 1 day after irradiation. Primary MEFs and HFFs were irradiated in triplicates in a 6-well plate, twice a day (acute exposure) and once every day for 3 consecutive days (chronic exposure). PBS solutions were collected immediately after the last UV treatments. After every treatment, the cells were washed with pre-warmed PBS and complete media was replenished for the accounted waiting timepoints. Thereafter, media solutions were collected 20 minutes and 24 hours after treatment cessation. These solutions were used to assess extracellular ROS signals. Cytosolic ROS production was evaluated after trypsinization of the cells, lysis of cellular membrane with 0.5% TritonX-100 and centrifugation for 5 minutes at 14,000 x g. Samples were loaded into a 96 black well plate, together with varying concentrations of hydrogen peroxide for generating the standard curve, and fluorescence signals were recorded using the Infinite 200 Tecan *i*-control plate reader machine at 480 nm excitation / 530 nm emission.

### Quantification of 8-hydroxy-2’-deoxyguanosine via ELISA

We quantified 8-hydroxy-2’-deoxyguanosine (8-oxo-dG) for oxidative DNA damage in DNA samples using the EpiQuik™ 8-OHdG DNA Damage Quantification Direct Kit (Colorimetric) from Epigentek following manufacturer instructions. Briefly, primary MEF, HFF, and HEKa primary cells were seeded on 10-cm dishes overnight. Three positive control exposures were employed, after concentration optimization, namely 400µM H_2_O_2_ for 1 hour, 20J/m^2^ UVC, and 4mM KBrO_3_ for 1 hour. Following irradiation with nail polish dryer, cells were collected at the selected timepoints (immediately, 4 hours, and 24 hours after irradiation) and subjected to genomic DNA extraction. Following Qubit DNA quantification, 300ng of DNA was loaded into strip wells that are specifically treated to have a high DNA affinity. 8-oxo-dG was detected using capture and detection antibodies. The detected signal is enhanced, and the readout was measured at 450 nm absorbance using the Infinite 200 Tecan *i*-control plate reader machine.

### Evaluation of mitochondrial ROS production and membrane potential

Image-iT™ TMRM Reagent, and MitoSOX™ Red Mitochondrial Superoxide Indicator (ThermoFisher Scientific; catalog number: I34361 and M36008, respectively), are two dyes that target mitochondria in live cells, the former localizing active mitochondria with normal mitochondria membrane potential, and the latter with affinity to ROS production. MitoTracker™ Green FM (ThermoFisher Scientific, catalog number: M7514) was utilized to localize mitochondria regardless of mitochondrial membrane potential. Briefly, primary cells were seeded on Mattek 35-mm dishes (P35G-1.5-14-C) overnight and incubated for 30 minutes in pre-warmed live imaging staining solution (ThermoFisher Scientific, catalog number: A14291DJ) containing 100µM MitoTracker reagent. TMRM dye was added in pre-warmed Live Imagining Solution to a final concentration of 100nM and incubated for 30 minutes at 37 °C. MitoSOX solution was prepared in pre-warmed Live Imagining Solution to a final concentration of 200nM and incubated for 10 minutes at 37 °C. All solutions were washed carefully 3 times with pre-warmed PBS and normal media was replenished. Live cell imaging was conducted using Nikon A1R-STORM Super Resolution Microscope and processed on Fiji software (version 2.3.0). Fluorescence data quantification was carried out using General Analysis pipeline on Nikon WorkStation 5317.

### DNA extraction and bulk whole-genome sequencing

Genomic DNA from MEF and HFF primary cells and immortalized clones were extracted using Qiagen DNeasy Blood & Tissue kit, following the manufacturer instructions. Quality and quantity of DNA were checked using NanoDrop and Qubit instruments. Around 2ug of DNA was thereby extracted per each sample. High-quality DNA for 32 samples were sent to Novogene for whole-genome library preparation and whole-genome sequencing at 30x coverage using paired-end 150 base-pair run mode with Illumina’s HiSeq-XTen. The 32 samples included: *(i)* one primary HFF and one primary MEF used as normal samples in the mutation calling; *(ii)* 5 HFF and 5 MEF immortalized clones after acute irradiation; *(iii)* 5 HFF and 5 MEF immortalized clones after chronic irradiation; and *(iv)* 5 HFF and 5 MEF unirradiated immortalized control clones.

### Identification of somatic mutations from whole-genome bulk sequencing

FASTQ files were subjected to BWA-MEM alignment using GRCm38 and GRCh38 as reference genomes for MEF and HFF, respectively. Our methodology for identification of somatic mutations from bulk sequencing data follows established approaches from large genomics consortia^83,84^. Briefly, ensemble variant calling of somatic mutations was performed using three independent variant callers Mutect2^85^, VarScan2^86^, Strelka2^87^, and MuSe^88^. Any mutation identified by at least two out of the four variant callers was considered a *bona fide* mutation. *Bona fide* mutations were subsequently filtered to remove any residual SNPs based on dbSNP annotation by variant effect predictor^89^. Further, any mutations shared between two or more samples and clustered mutations were removed as these reflect either residual germline mutations or mutations under positive selection (**Supplementary Figure 5**). Overall, a total of 118,429 unique somatic mutations were detected across all sequenced samples prior to filtering. Consistent with prior datasets^83,84^, the germline filtering removed 18.7% of these mutations with another 5.2% removed by the clustered filter. The remaining set of somatic mutations were used in the subsequent analyses and the evaluation for mutational signatures. Somatic copy-number changes were detected using FACETS^90^ with default parameters using the wrapper script cnv_facets v0.16.0.

### DNA extraction and duplex sequencing

Genomic DNA from HEKa cells were extracted using Qiagen DNeasy Blood & Tissue kit, following manufacturer instructions, with one exception during the initial proteinaseK digestion for which the samples were incubated at 37 rather than 54 for 1 hour, and DNA yield was eluted in IDTE buffer (10mM Tris HCl pH 8.0 + 0.1mM EDTA). Quality and quantity of DNA were checked using NanoDrop and Qubit instruments. High-quality DNA for 7 samples were sent to TwinStrand Biosciences for targeted library preparation, targeted duplex sequencing, and data analysis. The seven samples included: *(i)* one primary HEKa sample used in the mutation calling; 3 HEKa samples after chronic irradiation; and *(iii)* 3 HEKa unirradiated immortalized control clones. Briefly, duplex library preparation was performed using a human mutagenesis panel (∼50 kb distributed across the whole genome in regions not predicted to be under positive or negative selection) on DNA input sufficient to generate between 500 million to 1 billion informative Duplex base pairs per sample. Each sample was prepared in two batches: one with the standard mutagenesis assay and one with a prototype rapid workflow version of the assay. All libraries were sequenced on an Illumina NovaSeq 6000.

### Identification of somatic mutations from duplex sequencing

All raw NovaSeq sequencing FASTQ data was processed through the TwinStrand cloud-based human mutagenesis pipeline to generate error-corrected BAM files and variant call files for each sample. Duplex sequencing yielded an average of ∼239 million raw reads per sample, an average of ∼982 million duplex bases per sample, and an average of 15.204x on target duplex depth. No interspecies or intra-sample contamination found. Hybrid selection efficacy was approximately 99.81%. Mutagenesis analysis was carried out using the *Min* assumption for mutant frequency calculation as standardly done^39^. This method counts each variant only once, regardless of the number of reads that contain the non-reference allele. As previously done, the variant allelic frequency of each mutations was calculated by dividing the number of unique variants to the total number of duplex bases^39^. The set of somatic mutations with variant allelic frequency ≤ 1% were used in the subsequent analyses and the evaluation for mutational signatures.

### Analysis of mutational signatures and additional examinations

Analysis of mutational signatures was performed using our previously derived set of reference mutational signatures^59^ as well as our previously established methodology with the SigProfiler suite of tools used for summarization, simulation, visualization, and assignment of mutational signatures^91-94^. Variant allele frequencies (VAF) were calculated using integrative genomics viewer^95^. R version 3.6.1 was used to plot data (*i*.*e*., ggplot2, easyGgplot2, ComplexHeatmap^96^, and circlize^97^ packages), to compute p-values (ggpubr package), to perform correlation analyses (corrr package), and to compute the cosine similarity (lsa^98^ package).

## Statistics and Reproducibility

Unless otherwise annotated, all statistical comparisons were performed using Mann-Whitney U two-sided tests. All data generated in this study were collected from at least triplicates, unless otherwise stated. Each experiment was performed at least three times independently with reproducible results. All immunofluorescence images were collected from triplicates and each assay was performed three times. No statistical method was used to predetermine sample size. No data were excluded from the performed analyses. Our experiments were not randomized. The investigators were not blinded to allocations during experiments and outcome assessment.

## DATA AVAILABILITY

All whole-genome and duplex sequencing data have been deposited to Sequence Read Archive (SRA) and can be downloaded using accession number: PRJNA667106. All data and metadata for the previously generated whole-genome sequenced skin cancers were obtained from the official PCAWG release (https://dcc.icgc.org/releases/PCAWG). Where appropriate, source data are provided for the figures in the paper. For mouse samples, we aligned FASTQ file to the GRCm38 reference genome. For human samples, we aligned FASTQ files to the GRCh38 reference genome. We employed the dbSNP142 for germline mutations filtration in mouse samples, and dbSNP155 for human samples.

## CODE AVAILABILITY

Somatic mutations in whole-genome sequencing data were identified using our ensemble variant calling pipeline, which is freely available under the permissive 2-clause BSD license at: https://github.com/AlexandrovLab/EnsembleVariantCallingPipeline. All other computational tools utilized in this publication have been previously published and can be access through their respective publications.

## ACKNOWLEDGEMENTS

The authors would like to thank Dr. John Murray (Indiana University Bloomington) for providing human foreskin fibroblasts. This work was supported by an Alfred P. Sloan Research Fellowship to LBA. LBA is also an Abeloff V scholar, and he is personally supported by a Packard Fellowship for Science and Engineering. This work was also supported by the US National Institute of Health grants R01ES030993-01A1, R01ES032547-01, and R01CA269919-01 to LBA as well as R35ES031638-03 to BVH. The funders had no roles in study design, data collection and analysis, decision to publish, or preparation of the manuscript.

## AUTHOR CONTRIBUTIONS

LBA and MZ designed all experiments with help from BVH. MZ performed all experiments with assistance from AH, NV, SPN and YY. MZ performed all data analysis with help from CY, JD, BO, and YH. BVH assisted in the writing of the manuscript and helped interpreting the experimental results. The manuscript was written by LBA and MZ with input from all other authors. LBA supervised the overall project and writing of the manuscript. All authors read and approved the final manuscript.

