## Supplementary Information for "DNA damage and somatic mutations in mammalian cells after irradiation with a nail polish dryer"

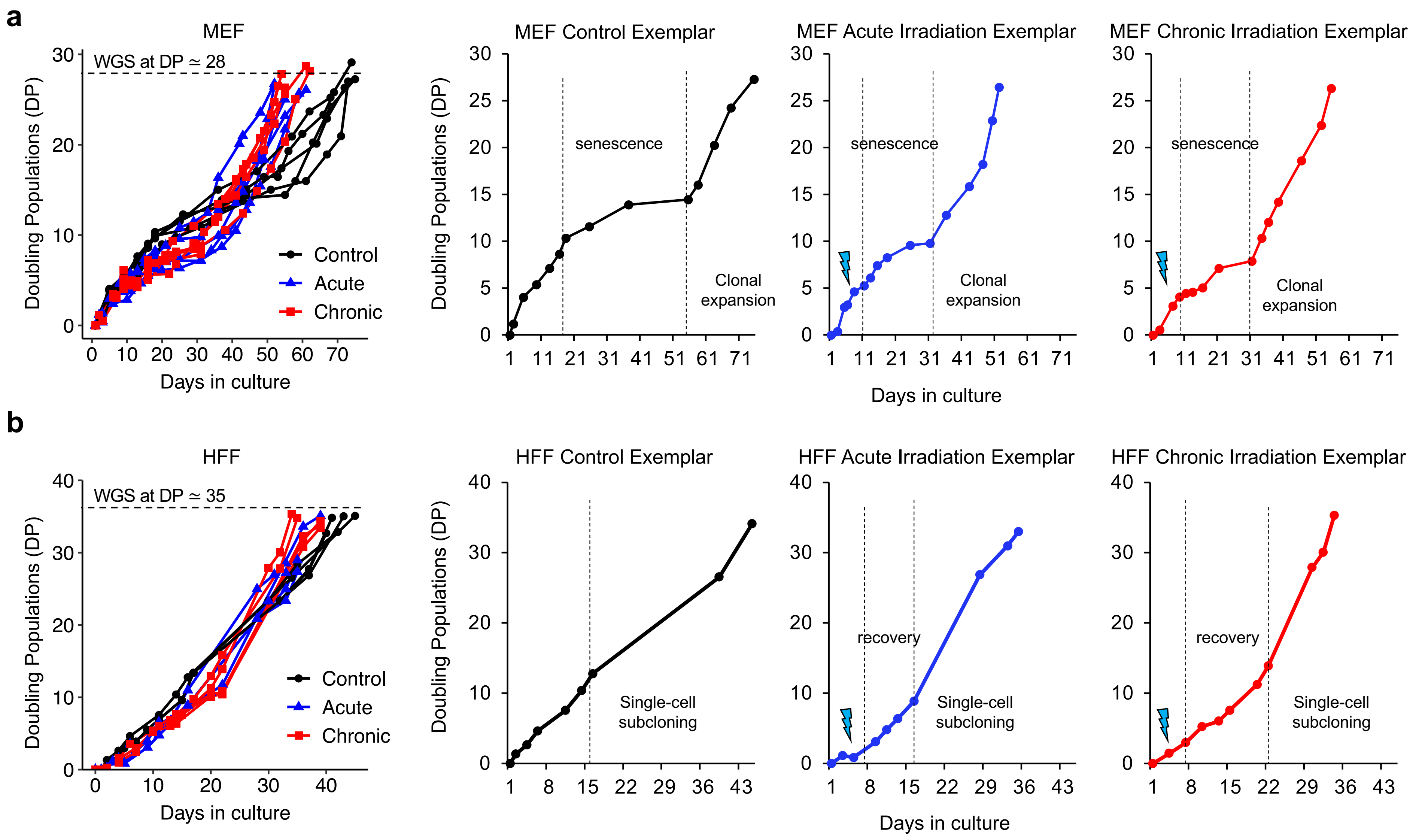

**Supplementary Figure 1. Exposure and barrier bypass-clonal expansion assays.** ***(a)*** MEFs were treated at early passages and led through senescence manifested by a plateau-like curve where cells lost the ability to efficiently duplicate and grow. Cells went through senescence bypass and eventually emerge into immortalized clones, able to multiply indefinitely, a process called barrier-bypass clonal expansion, emulating oncogenesis. Irradiated and control MEF cells were grown for ~28 doubling populations prior to bulk whole-genome sequencing (WGS). Exemplars are provided for one control, one acutely irradiated, and one chronically irradiated MEF. Y-axes reflect the number of doubling populations, while X-axes correspond to the number of days in culture. ***(b)*** HFFs were treated at early passages and led through recovery phase before initiating a single cell subcloning assay using a serial dilutions technic. *Bona fide* single-cell clones were collected at the end of experiment. Irradiated and control HEF cells were grown for ~35 doubling populations prior to bulk whole-genome sequencing (WGS). Exemplars are provided for one control, one acutely irradiated, and one chronically irradiated HFF. Y-axes reflect the number of doubling populations, while X-axes correspond to the number of days in culture.

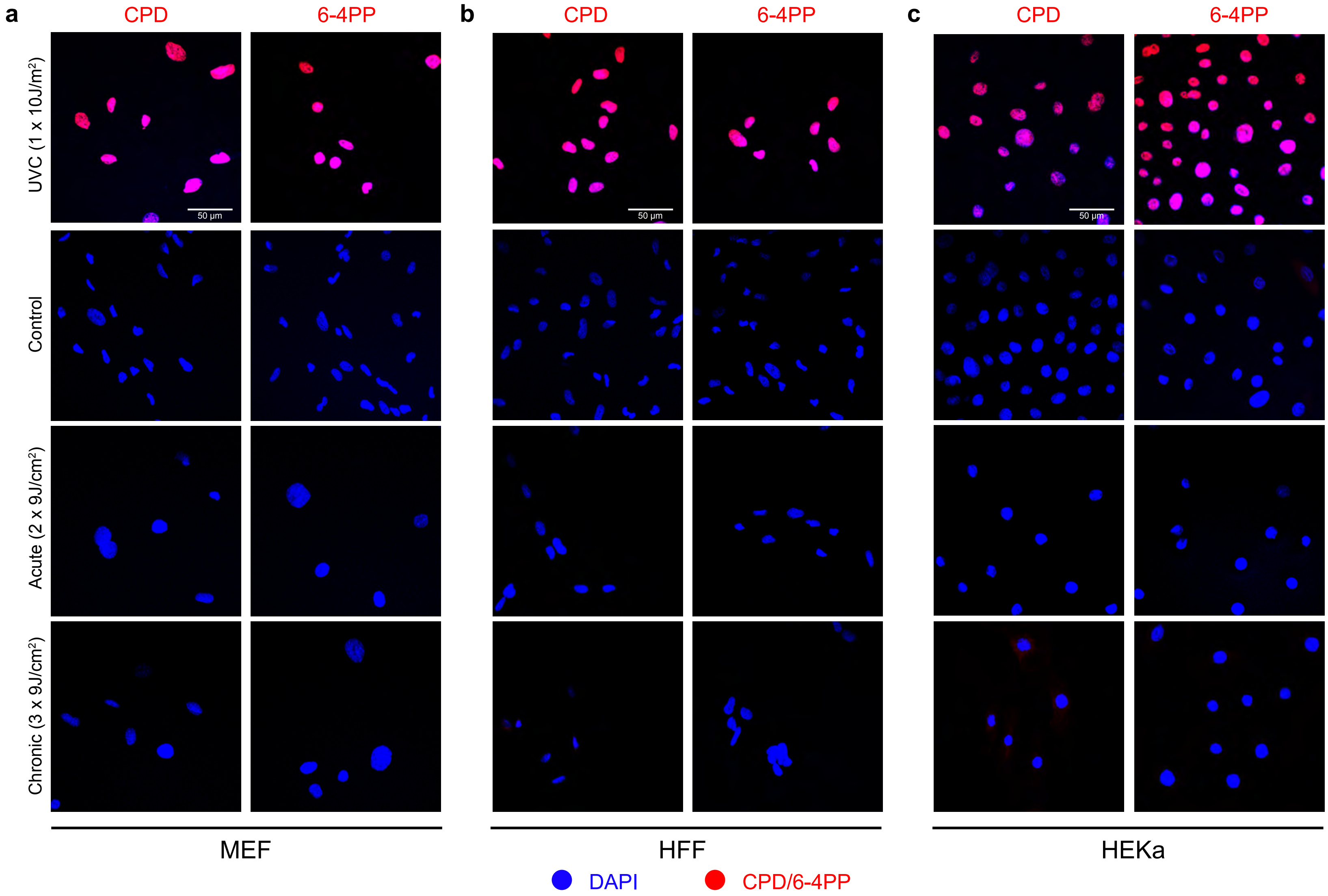

**Supplementary Figure 2. Evaluation of cyclobutane pyrimidine dimers and 6-4 photoproducts irradiated by a UV-nail dryer.** ***(a-c)*** Immunofluorescence of cyclobutane pyrimidine dimers (CPDs) and 6-4 photoproducts (6-4PPs) in MEF ***(a)***, HFF ***(b)***, and HEKa ***(c)*** cells after acute or chronic irradiation with a UV-nail dryer. DAPI is shown in blue while CPDs and 6-4PPs are shown in red. A positive control, namely, cells irradiated with ultraviolet C (UVC) light (1x 10J/$m^2$), is also provided for each cell line. Images collected from four independent experiments for MEFs and HFFs and five independent experiments for HEKa cells.

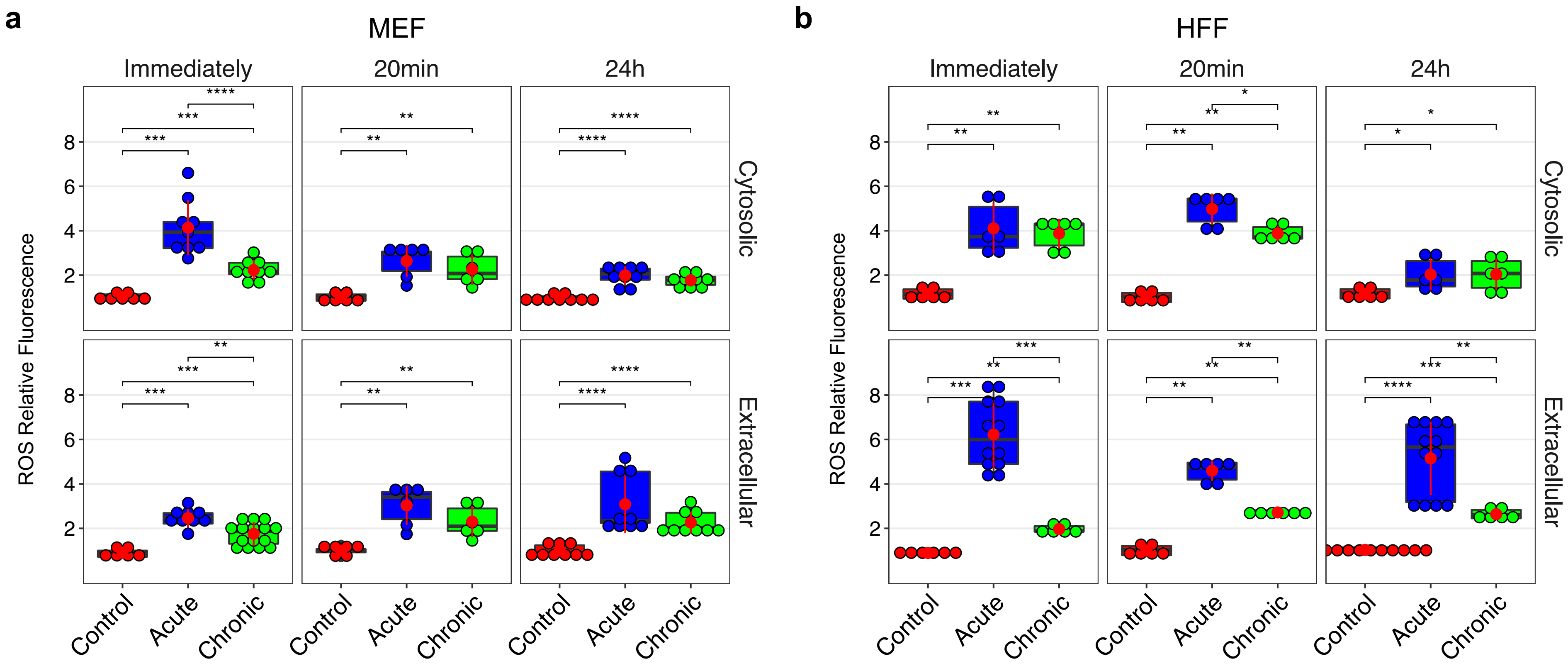

**Supplementary Figure 3. Cytosolic and extra-cellular measurement of reactive oxygen species in MEFs and HFFs.** Accumulation of reactive oxygen species (ROS) through time in primary MEFs ***(a)*** and HFFs ***(b)*** after acute or chronic irradiation with a UV-nail dryer. Each circle corresponds to a single experiment for a particular exposure. All y-axes reflect, either cytosolic or extracellular, relative ROS fluorescence by normalizing the fluorescence of a particular exposure to the average fluorescence of the appropriate controls. Data collected from $n=9$ independent experiments for MEFs (exceptions apply for all ROS measurements 20 minutes after exposure where $n=6$, and for ROS measurement extracellularly immediately after chronic exposure in MEFs and in control samples where $n=15$) and $n=6$ independent experiments for HFFs (exception for ROS measurement extracellularly immediately and 24h after acute irradiation in HFFs and in control samples at 24h where the used number of independent biological replicates is $n=12$). The bounds of the boxplots represent the interquartile range divided by the median, and Tukey-style whiskers extend to a maximum of 1.5 × interquartile range beyond the box. Statistically significant results from FDR corrected Mann-Whitney U two-sided tests are denoted as: **q-value<0.01, ***q-value<0.001 and ****q-value<0.0001.

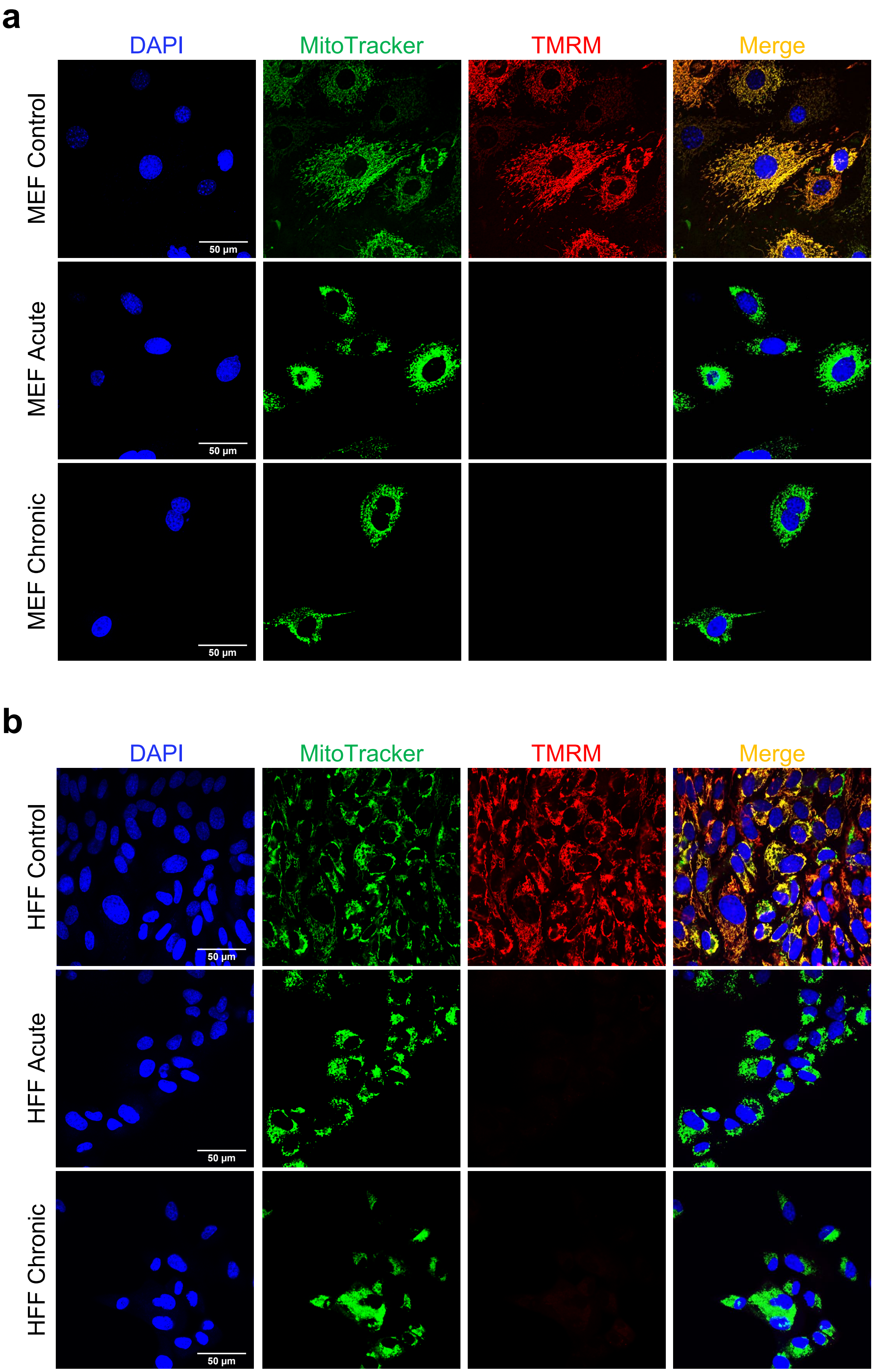

**Supplementary Figure 4. Mitochondrial membrane potential after irradiation with a UV-nail dryer.** Evaluating the mitochondrial membrane potential after irradiation of MEF cells ***(a)*** and HFF cells ***(b)*** with a UV-nail dryer. DAPI is shown in blue, MitoTracker in green, and tetramethylrhodamine dye (TMRM) in red. Yellow corresponds to overlaps between MitoTracker and TMRM. Images collected from three independent experiments for MEFs and HFFs.

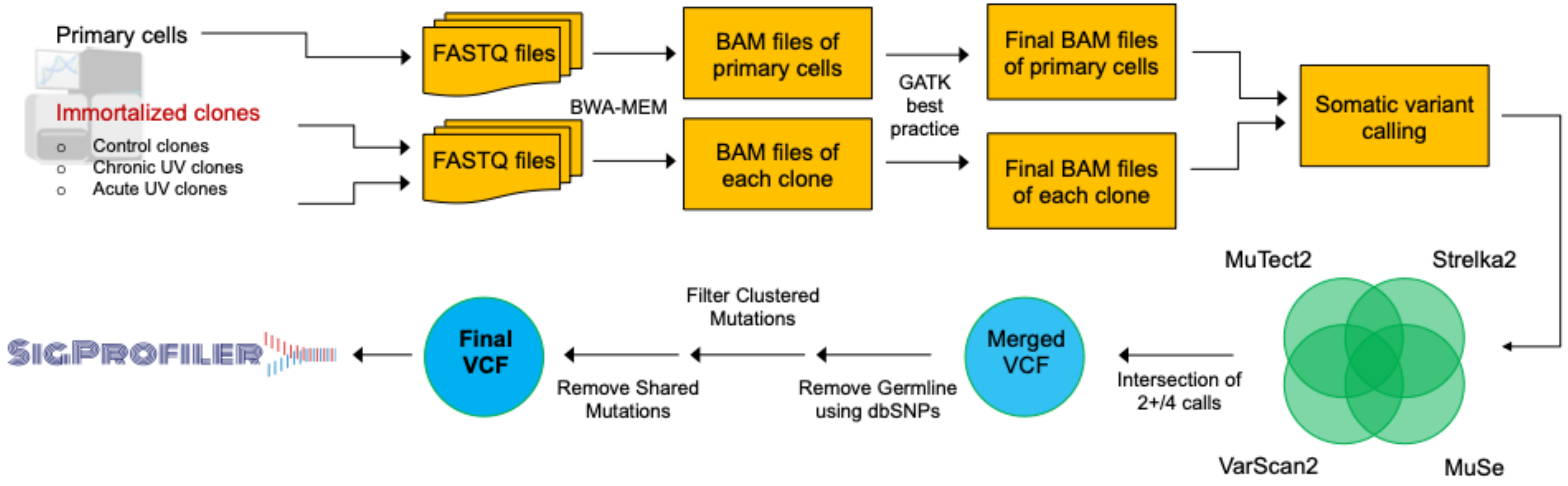

**Supplementary Figure 5. Schematic of whole-genome data analysis.** Primary cells were sequenced and used as a germline reference. We employed three variant callers (Mutect2, VarScan2, Strelka2, and MuSe) in matched tumor-normal mode, with the primary cells' BAM files as normal for each cell line. We selected variants called in at least 2 of the variant callers in order to increase variant calling quality. Germline single nucleotide polymorphisms (SNP) were removed using the dbSNP database. Any clustered mutation as well as any mutation observed in 2 or more samples were also removed. The final set of somatic mutations were analyzed by the SigProfiler suite of tools.

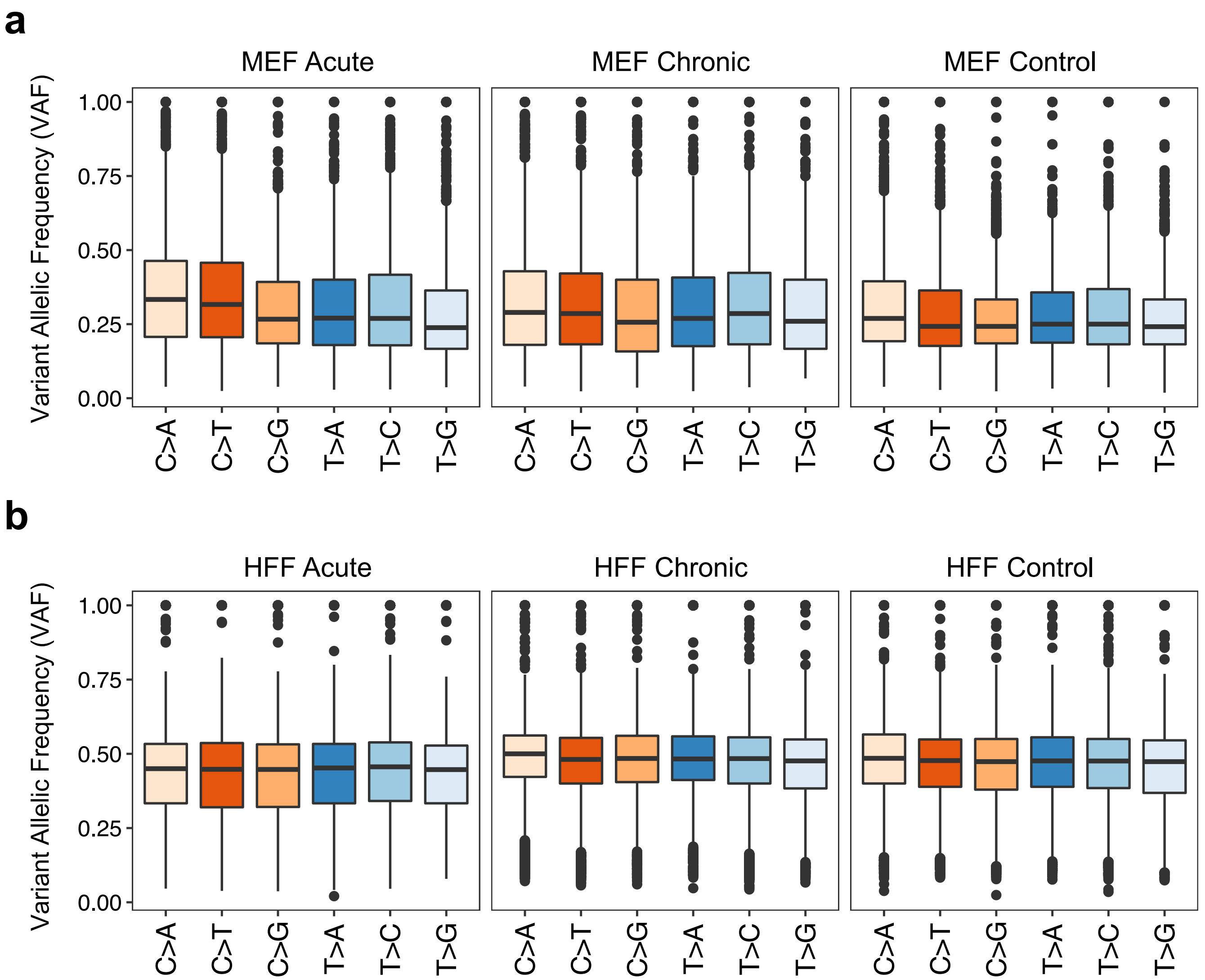

**Supplementary Figure 6. Analysis of variant allelic frequency analysis.** Variant allelic frequency of *bona fide* mutations in MEF clones ***(a)*** HFF clones ***(b)*** for each irradiation condition. Data collected from *n=5* independent biological replicates for each condition in MEFs and HFFs. The bounds of the boxplots represent the interquartile range divided by the median, and Tukey-style whiskers extend to a maximum of 1.5 × interquartile range beyond the box.

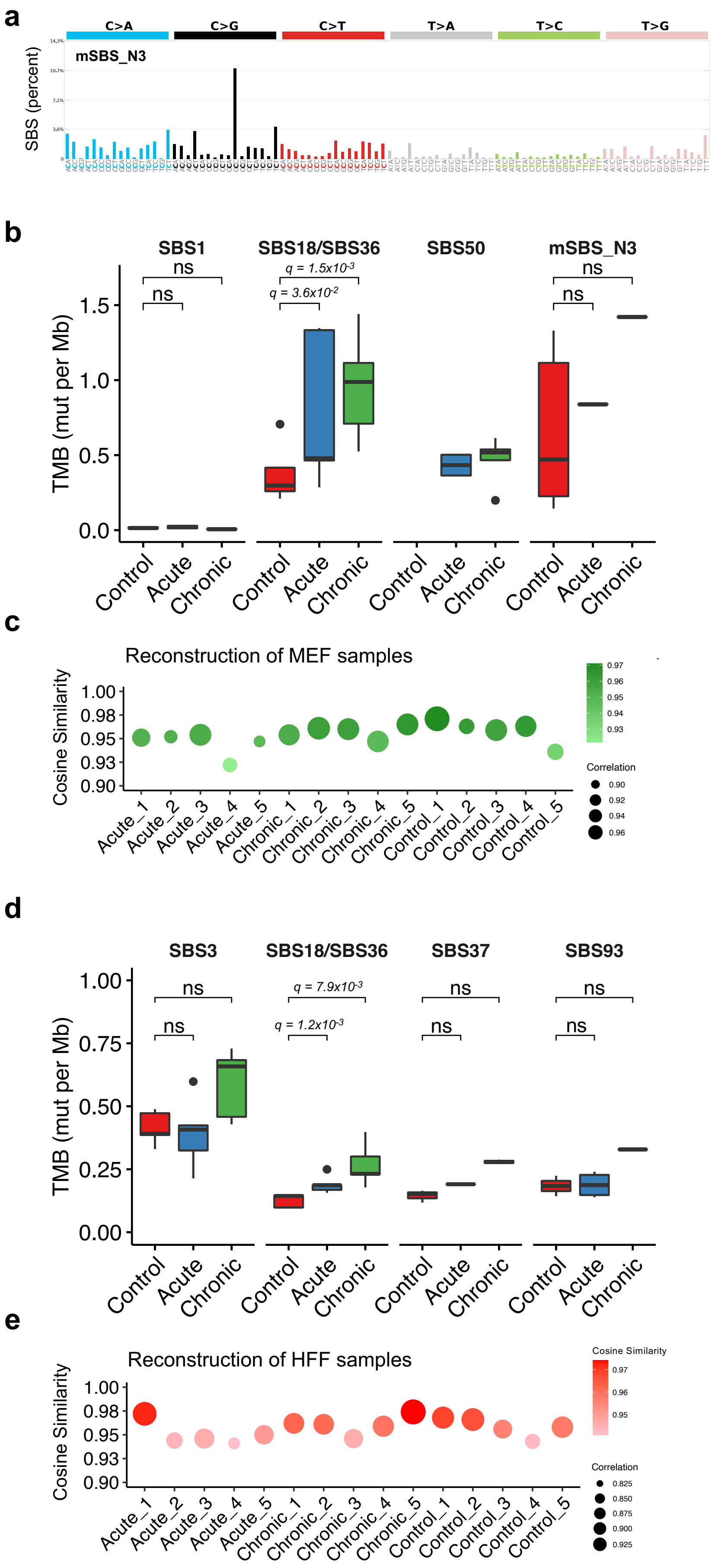

**Supplementary Figure 7. Mutation analysis of UV-treated MEF and HFF clones and PCAWG samples.** ***(a)*** Mutational signature mSBS_N3 identified *in vivo* in mouse models. ***(b)*** COSMIC signatures assigned to MEF clones per condition (control, acute and chronic). Data collected from *n=5* independent biological replicates for each condition. Statistically significant results from FDR corrected Mann-Whitney U two-sided tests are denoted as *q*-values. **(*c*)** Mutational signatures reconstruction of MEF mutational patterns. Accuracy is measured in cosine similarity between the original pattern of a sample and the pattern of a sample reconstructed using the assigned mutational signatures. **(*d*)** COSMIC signatures assigned to HFF clones per condition (control, acute and chronic). Data collected from *n=5* independent biological replicates for each condition. Statistically significant results from FDR corrected Mann-Whitney U two-sided tests are denoted as *q*-values. **(*e*)** Mutational signatures reconstruction of HFF mutational patterns. The bounds of the boxplots represent the interquartile range divided by the median, and Tukey-style whiskers extend to a maximum of 1.5 × interquartile range beyond the box.

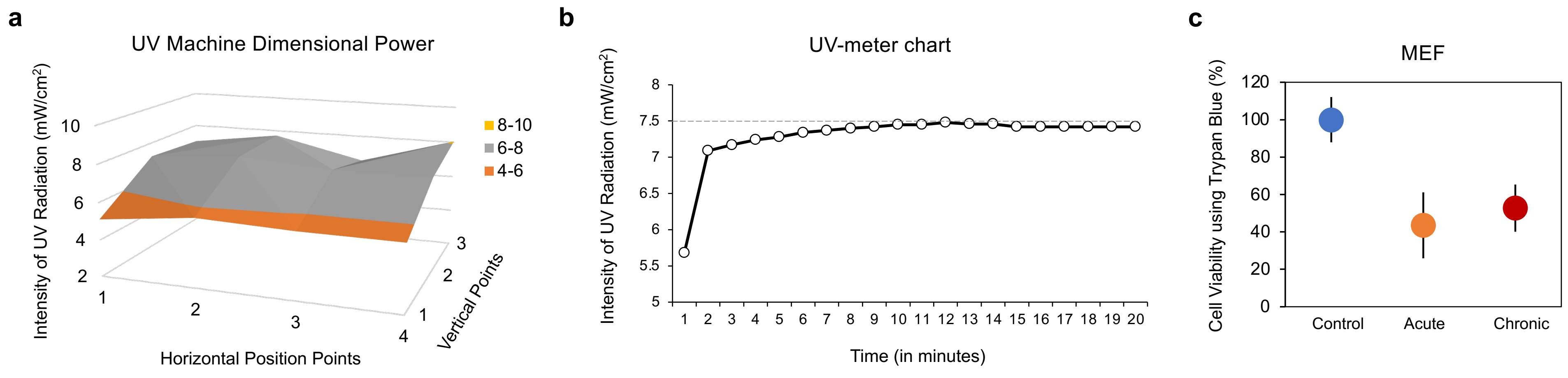

**Supplementary Figure 8. Toxicity and characteristics of the UV-nail polish dryer machine.** ***(a)*** Intensity of UV radiation (mW/cm$^2$) at different position points (vertical and horizontal) in the employed UV nail polish dryer. ***(b)*** Stability assessment of the UV intensity emitted from the UV-nail polish dryer over time. ***(c)*** Validation of the half maximal inhibitory concentration (IC50) used for the acute and chronic exposure protocols in MEFs. Data presented as mean viability percent ± standard deviation from *n=3* independent biological replicates.

### SUPPLEMENTARY TABLES

**Supplementary Table 1. Assays and timepoints for assessing DNA damage.** The table denotes timepoints at which DNA damage and other experimental assessments were performed for each condition in primary cell models, including interrogation of cytotoxicity, genotoxicity, oxidative damage, and mitochondrial damage.

| Assessment | Assays | Timepoints |
|---|---|---|
| Cytotoxicity | Trypan Blue | 48 hours |
| Cytotoxicity | Cell Counting kit -8 | 49 hours |
| Genotoxicity | ɣH2Ax immunofluorescence | 4 hours |
| Genotoxicity | CPD/6-4PP immunofluorescence | Immediately |
| Oxidative Damage | CellROX green reagent | Immediately |
| Oxidative Damage | OxiSelect In vitro ROS/RNS assay kit | Immediately |
| Oxidative Damage | OxiSelect In vitro ROS/RNS assay kit | 20 minutes |
| Oxidative Damage | OxiSelect In vitro ROS/RNS assay kit | 24 hours |
| Oxidative Damage | EpiQuik 8-OHdG DNA damage quantification kit | Immediately |
| Oxidative Damage | EpiQuik 8-OHdG DNA damage quantification kit | 4 hours |
| Oxidative Damage | EpiQuik 8-OHdG DNA damage quantification kit | 24 hours |
| Mitochondrial Damage | TMRM mitochondrial membrane potential indicator | 24 hours |
| Mitochondrial Damage | MitoSOX mitochondrial superoxide indicator | 25 hours |
